# Distinct and Cooperative Roles of DNA Methylation and Meiotic Chromosome Architecture in Crossover Control

**DOI:** 10.64898/2026.08.04.742853

**Authors:** Chiara Di Dio, Denitsa Hrisova, Natasha E. Yelina

## Abstract

During meiosis, homologous chromosomes exchange segments in a process termed crossover recombination. Crossovers are non-randomly distributed along chromosomes, and in many eukaryotes, including plants, meiotic chromosome architecture and chromatin states control recombination landscapes. Whether these two components genetically interact has remained underexplored.

To address this question, we combined *Arabidopsis thaliana*, hereinafter, Arabidopsis, mutations that disrupt meiotic chromosome architecture by depleting the meiotic chromosome axis (*asy1/+*) or synaptonemal complex (*zyp1*) with mutations in the DNA methyltransferases *MET1* and *CMT3* (*met1/+* and *cmt3*), which lead to a loss of cytosine DNA methylation, the hallmark of heterochromatin, in the CG and CHG contexts, respectively. We quantified crossovers in telomere- and centromere-proximal chromosome intervals using fluorescent seed-based reporters and found that DNA methylation and meiotic chromosome architecture proteins can have distinct or cooperative roles in crossover control depending on the chromosome interval and DNA methylation context. We demonstrate that axis and synaptonemal complex act together with CG DNA methylation to control crossovers, while CHG DNA hypomethylation cannot fully restore a loss of centromere-proximal recombination caused by the depletion of *ASY1* or *ZYP1*.

Remarkably, increasing *ASY1* dosage promotes crossovers within the pericentromere, representing a new non-epigenetic route to upregulate pericentromeric recombination.

**Author summary:** Meiotic crossovers reshuffle genetic variation and are essential for evolution and crop breeding. However, crossovers occur unevenly along chromosomes, limiting genetic exchange in pericentromeric regions. Here, we investigate the genetic interactions between cytosine DNA methylation and meiotic chromosome architecture and show that, although heterochromatin depletion can permit pericentromeric crossovers, the structural integrity of the meiotic chromosome axis and the synaptonemal complex are essential to drive recombination. Remarkably, modulating the dosage of a chromosome axis protein provides a non-epigenetic strategy to increase pericentromeric crossovers, revealing new opportunities to reshape recombination landscapes in model and crop plants.

## Introduction

Meiosis is a specialized cell division where one round of DNA replication is followed by two consecutive cell divisions to produce haploid gametes required for sexual reproduction (Villeneuve and Hillers 2001; Mercier et al. 2015). During meiotic prophase I, homologous chromosomes undergo programmed recombination events that can result in reciprocal exchanges of genetic material, known as crossovers (Villeneuve and Hillers 2001; Mercier et al. 2015). Crossovers contribute to genetic variation in progeny, generating new haplotypes that can bring together agronomically valuable traits in crop species (Taagen et al. 2020). However, crossover recombination frequencies and distribution along chromosomes can limit breeding, as crossovers are relatively rare (typically 1–2 per chromosome per meiosis) and unevenly distributed along chromosomes (Mercier et al. 2015; Taagen et al. 2020). For example, crossovers in wheat, barley, soybean and maize occur predominantly in sub-telomeric regions (Higgins et al. 2012; Rogers-Melnik et al. 2015; Darrier et al. 2017; Mascher et al. 2017; Ma et al. 2023). This suppression of recombination at centromeres and centromere-proximal regions (hereinafter referred to as pericentromeres) frequently results in linkage drag. Therefore, technologies capable of increasing crossover frequencies genome-wide or within crossover-suppressed chromosomal regions have the potential to substantially accelerate crop breeding.

Crossovers are initiated by DNA double-strand breaks (DSBs) catalyzed by the transesterase SPO11 (Keeney et al. 1997; Bergerat et al. 1997). DSBs are processed to generate 3’ single-strand DNA (ssDNA) overhangs which associate with the recombinases RAD51 and DMC1 to promote ssDNA invasion of a homologous chromosome or a sister chromatid (Hunter 2015). Invasion of homologous DNA generates a displacement loop (D-loop), which can either be dissociated to yield a noncrossover, or progress to form a double Holliday junction (dHJ) joint molecule (Hunter 2015). In plants, noncrossover repair is actively promoted by FANCM, RECQ4A and RECQ4B, TOPOISOMERASE3α, FIGL1, and FLIP1 (Crismani et al. 2012; Girard et al. 2015; Séguéla-Arnaud et al. 2015; Fernandes et al. 2018). Alternatively, the Class I crossover pathway acts to stabilize dHJs and promote their resolution as crossovers. Class I crossovers make up to ∼85% of Arabidopsis crossovers and constitute an interfering pathway, where crossover events are more widely spaced out than expected by chance (Mercier et al. 2015). The remaining ∼15% minority of crossovers are noninterfering (Mercier et al. 2015). Overall, in Arabidopsis, ∼250 DSBs mature into only ∼10 crossovers per meiosis, while the remaining DSBs are repaired as noncrossovers (Mercier et al. 2015).

Homologous recombination depends on the formation of the chromosome axis, a dynamic, meiosis-specific protein assembly built along the DNA. Following the DNA synthesis (S) phase, replicated sister chromatids associate with cohesin complexes containing the meiosis-specific REC8 which is believed to serve as a structural foundation for meiotic chromosome axis assembly (Bhatt et al. 1999; Chelysheva et al. 2005; Golubovskaya et al. 2006; Shao et al. 2011). Other components of the meiotic chromosome axis include the coiled-coil proteins ASY3 (homolog of yeast Red1 and mammalian SYCP2) and ASY4 (homolog of mammalian SYCP3), alongside the HORMA domain protein ASY1 (homolog of yeast Hop1 and mammalian HORMAD1/2) (Ross et al. 1997; Armstrong et al. 2002; Wojtasz et al. 2009; Ferdous et al. 2012; Chambon et al. 2018). This organization establishes an axis-loop structure where chromatin loops project laterally from the core axis (Zickler and Kleckner 1999). Axis-localized ASY1 serves as a key coordinator of meiotic recombination, ensuring chromosome pairing, crossover interference and assurance, and genome stability. Consequently, the loss of ASY1 leads to a loss of crossover interference, a dramatic reduction in genome-wide crossover numbers, disrupted meiotic fidelity, chromoanagenesis, and sterility (Shin et al. 2010; Daniel et al. 2011; Mercier et al. 2015; Lambing et al. 2020; Guo et al. 2023; Pochon et al. 2023; Wang et al. 2023). Mechanistically, ASY1 directly recruits plant-specific cyclin SDS to chromosomes, which promotes efficient DMC1 localization and interhomolog recombination (Cai et al. 2026). The precise regulatory impact of ASY1 varies among species: in Arabidopsis, ASY1 acts in a dosage-dependent manner and is believed to antagonize telomere-led recombination (Lambing et al. 2020). Conversely, in barley, chromosome axis formation is initially polarized to distal regions mirroring its subtelomeric crossover distribution (Higgins et al. 2012). In polyploid crops like wheat, in addition to maintaining crossover assurance and interference, ASY1 suppresses illegitimate recombination between non-homologous chromosomes (Di Dio et al. 2023).

Once homologs pair and synapse, the chromosome axis is modified as homologous chromosomes are held together by a proteinaceous structure called the synaptonemal complex (SC) (Zickler and Kleckner 2015; Page and Hawley, 2004). The SC features lateral elements connected by a central region composed of the transverse filament protein ZYP1 (homolog of yeast Zip1 and mammalian SYCP1) (Sym et al. 1993; Higgins et al. 2005; De Vries et al. 2005; Qiao et al. 2012; Barakate et al. 2014). In *Arabidopsis*, ZYP1 is not required for crossover formation itself, but is essential for maintaining crossover interference and underpins heterochiasmy, or sex-specific differences in crossover landscapes (Capilla-Pérez et al. 2021; France et al. 2021).

Chromatin is another critical determinant of crossover control. Plant heterochromatin is epigenetically modified with DNA cytosine methylation and histone H3K9me2 methylation, which suppress the transcriptional activity of transposable elements. DNA methylation occurs in CG, CHG, and CHH contexts (where H = A, C, or T) (Lippman et al. 2004; Du et al. 2012; Stroud et al. 2013; Stroud et al. 2014). CG methylation is maintained by METHYLTRANSFERASE1 (MET1), while non-CG methylation is maintained redundantly by the cytosine methyltransferases CHROMOMETHYLASE2 (CMT2), CHROMOMETHYLASE3 (CMT3), and DOMAINS REARRANGED METHYLASE2 (DRM2) (Bartee et al. 2001; Lindroth et al. 2001; Saze et al. 2003; Du et al. 2012; Stroud et al. 2013; Stroud et al. 2014). Consequently, plant crossovers are excluded from heterochromatin-dense centromere-proximal regions and predominantly occur at gene transcription start and termination sites characterized by low nucleosome density and active histone marks (H3K4me3 and H2A.Z) (Choi et al. 2013; Hellsten et al. 2013; Choi and Henderson 2015). Targeted recruitment of heterochromatin actively suppresses crossover formation (Yelina et al. 2015).

Consistent with the excess of meiotic DSBs over crossovers, plant crossovers are below their potential maximum. In line with this, multiple routes have been developed to increase crossover numbers and alter crossover landscapes in plant chromosomes. Thus, knocking down anti-crossover factors (such as FANCM, RECQ4A, and RECQ4B helicases, TOPOISOMERASE3α, FIGL1, and FLIP1) (Crismani et al. 2012; Girard et al. 2015; Séguéla-Arnaud et al. 2015; Fernandes et al. 2018), depleting ZYP1 (Capilla-Pérez et al. 2021), upregulating the dosage-acting Class I pro-crossover protein HEI10 (Ziolkowski et al. 2017), or a combination of these approaches leads to global crossover increases (Serra et al. 2018; Durand et al. 2022; Jing et al. 2025). Knocking out cohesin regulators, SGO2 and CTF18, as well as deSUMOylase SPF2, upregulates pericentromeric crossovers (Salinas Gamboa et al. 2026). Depletion of CG DNA methylation in *met1* mutants or reducing ASY1 dosage remodels crossovers from pericentromeres to sub-telomeres (Yelina et al. 2012; Yelina et al. 2015; Lambing et al. 2020), while depletion of CHG DNA methylation and H3K9me2 in *cmt3* mutants results in epigenetic activation of recombination near Arabidopsis centromeres (Underwood et al. 2018).

Despite this wealth of knowledge, whether the meiotic chromosome axis and synaptonemal complex, which we collectively refer to as ‘meiotic chromosome architecture proteins’, genetically interact with heterochromatin to control crossovers remains underexplored. Here, we address this question by combining mutants that lead to CG and CHG DNA hypomethylation with mutants depleted for the meiotic chromosome axis and synaptonemal complex components. We find evidence for both, cooperative and distinct functions of meiotic chromosome architecture and cytosine DNA methylation in controlling pericentromeric and sub-telomeric crossovers. We show that while CHG DNA hypomethylation is insufficient to compensate for a centromere-proximal crossover loss caused by ASY1 or ZYP1 depletion, increasing ASY1 dosage leads to an upregulation of centromere-proximal crossovers, identifying a new non-epigenetic approach to activate recombination in plant pericentromeres.

## Materials and Methods

### Plant material and genotyping

Arabidopsis lines used in this study were Col-0, *asy1-4* (SALK_046272, Col-0 accession), *cmt3-11* (SALK_148381, Col-0 accession) (Chan et al. 2006), *met1-3* (Col-0 accession) (Saze et al. 2003), *CTL3.9* and *CTL3.2* (Col-0 accession) (Wu et al. 2015). The lines were obtained from the Eurasian Arabidopsis Stock Centre (uNASC) and Arabidopsis Biological Resource Centre (ABRC). *zyp1a zyp1b* (France et al. 2021) was a kind gift from Prof James Higgins (University of Leicester). Plants were grown at 19°C and 65% humidity with a long day photoperiod (16 h light) and light intensity of 250 μmol. PCR genotyping of *asy1-4*, *zyp1a zyp1b*, *cmt3-11*, and *met1-3* was performed using oligonucleotides listed in **Supplemental Table 1**.

### Generating ASY1-overexpressing (*ASY1-OE*) lines

To generate *ASY1-OE* lines, *ASY1* genomic region including its native promoter and terminator identified in a previous study (Yang et al. 2020) was amplified as two fragments with oligonucleotides listed in **Supplemental Table 1** and cloned via Golden Gate cloning using PaqCI restriction enzyme (New England Biolabs) into a pTRANS_210d binary vector, a kind gift from Dr Evan Ellison (Crop Science Centre, University of Cambridge, and Donald Danforth Plant Science Centre, USA). Golden Gate cloning design was performed in Benchling (https://benchling.com); cloning was performed following the New England Biolabs’ protocol for PaqCI Golden Gate assembly.

*pTRANS_210d-ASY1* constructs were verified by Sanger sequencing; Arabidopsis transformation was performed via floral dipping (Zhang et al. 2006). T_1_ transformants were selected on Murashige-Skoog media supplemented with vitamins, 0.8% agar and 15 µg/ml hygromycin B following an established protocol (Harrison et al. 2006).

### Crossover quantification

Crossover quantification in *CTL3.2* and *CTL3.9* intervals was performed as described previously (Carpenter et al. 2006; Kbiri et al. 2022; Yelina et al. 2022).

### RT-qPCR

Closed flower buds from 3 inflorescences on an individual plant were pooled and disrupted in liquid N_2_. RNA was extracted using RNeasy Plant Mini Kit (Qiagen) following the manufacturer’s protocol. 2.5 µg RNA per sample was DNase treated using DNase I (Thermo Fisher Scientific), following the manufacturer’s protocol. Reverse transcription was performed using SuperScript IV reverse transcriptase (Thermo Fisher Scientific), following the manufacturer’s protocol. Quantitative PCR (qPCR) was performed using Luna Universal qPCR Master Mix (New England Biolabs), following the manufacturer’s protocol. Meiotic *DMC1* was used as a housekeeping control gene. Oligonucleotides used for qPCR are listed in **Supplemental Table 1**.

## Results

### CG DNA hypomethylation and depletion of the synaptonemal complex transverse filament genetically interact to control sub-telomeric and pericentromeric crossovers

To investigate whether CG DNA methylation and the synaptonemal complex genetically interact to control meiotic crossovers, we combined *zyp1a zyp1b* double mutants (hereinafter, *zyp1*), which lack the transverse filament, a key component of the synaptonemal complex (Capilla-Pérez et al. 2021; France et al. 2021), with *met1/+* heterozygotes, which exhibit CG DNA hypomethylation due to a mutation in *METHYLTRANSFERASE1* (*MET1*), an enzyme responsible for CG DNA methylation maintenance (Saze et al. 2003). We used *met1/+* heterozygotes rather than *met1* knockouts because, as shown previously, *met1/+* phenocopies the genome-wide crossover profiles of *met1* without causing stochastic silencing of the fluorescent reporters used to quantify crossovers (Yelina et al. 2015).

Here, we used fluorescent seed-based reporters to measure crossover frequencies in two contrasting intervals on chromosome 3 (**Figure 1A**): the heterochromatic centromere-proximal (hereinafter, pericentromeric) interval *CTL3.9* (9,741,508 - 15,980,483 bp) and the euchromatic sub-telomeric interval *CTL3.2* (130,439 - 4,330,342 bp) (Wu et al. 2015). These seed-based reporters represent genetically linked transgenes encoding *eGFP* and *dsRed* under the control of the seed-specific *napA* storage protein promoter (Wu et al. 2015). A crossover occurring within a reporter interval results in segregation of the *eGFP* and *dsRed* markers (**Figure 1B-D**), allowing robust quantification of crossover recombination frequencies within the reporter intervals. *CTL3.9* and *CTL3.2* reporters were introgressed into *zyp1*, *met1/+* and *zyp1 met1/+* via crossing to measure recombination frequencies.

**Figure 1.**
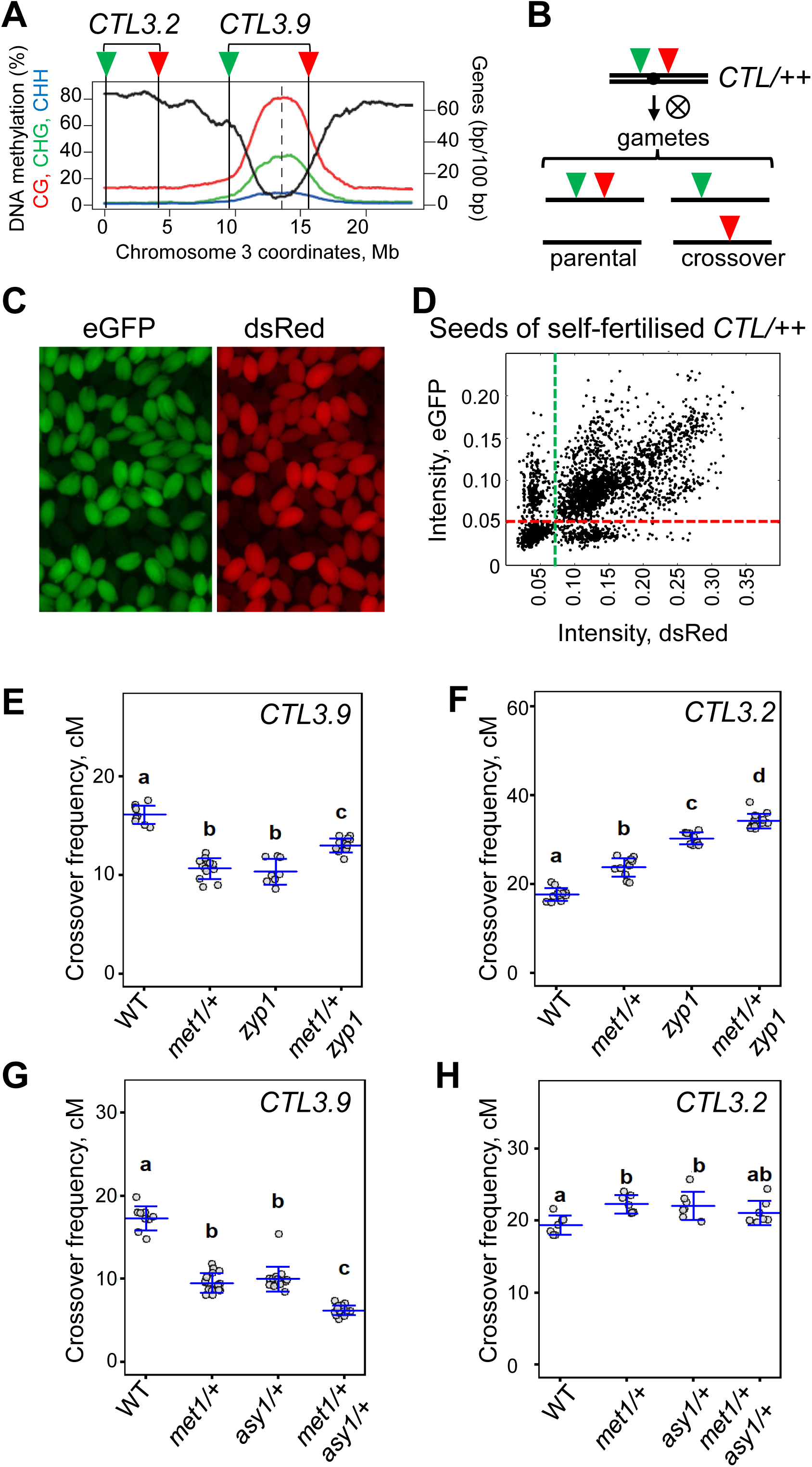
Genetic interactions between CG DNA hypomethylation and meiotic chromosome architecture in crossover control. **(A)** CG, CHG and CHH DNA methylation (red, green and blue, respectively, percentage) and gene density (black) in wild type Col-0 Arabidopsis chromosome 3. Centromere is indicated by a vertical dashed line. *eGFP* and *dsRed* of *CTL3.2* and *CTL3.9* reporters are indicated by green and red triangles, respectively, and vertical black lines. **(B)** Gametes produced by a plant carrying a hemizygous *CTL* reporter in the absence or presence of a crossover. **(C)** Fluorescent micrographs showing *CTL3.9/++* (*eGFP dsRed/++*) seed using green or red fluorescent filters. **(D)** A scatter plot of *eGFP* and *dsRed* fluorescence intensities in seeds produced by a plant carrying a hemizygous *CTL* reporter. Seeds showing fluorescence in either *eGFP,* or *dsRed* or neither *eGFP* or *dsRed* are separated by dashed red and green lines. **(E)** Crossover frequencies (cM) in *CTL3.9* in wild type, *met1/+*, *zyp1*, *met1/+ zyp1*. Measurements from individual plants are shown as circles. Horizontal blue lines indicate means and standard deviation. Different letters indicate statistically significant differences between groups determined by a global Kruskal-Wallis test followed by pairwise Wilcoxon tests with a Holm adjustment. **(F)** As in (E) but for *CTL3.2.* **(G)** As in (E) but for wild type, *met1/+*, *asy1*, *met1/+ asy1*. **(H)** As in (G) but for *CTL3.2*.

We found that, in agreement with the previous reports (Mirouze et al. 2012; Yelina et al. 2012; Yelina et al. 2015; Capilla-Pérez et al. 2021), crossovers in the pericentromeric *CTL3.9* interval were reduced in single *zyp1* and *met1/+* compared to the wild type (Wilcoxon test, *p*-value < 0.01, **Figure 1E**, **Supplemental Tables 2, 3>**). We found that combining *zyp1* with *met1/+* led to a modest, but statistically significant pericentromeric crossover increase compared to *zyp1* (Wilcoxon test, *p*-value = 1.6*10^−4^) or *met1/+* alone (Wilcoxon test, *p*-value = 1.3*10^−3^, **Figure 1E**, **Supplemental Tables 2, 3>**), suggesting that combined depletion of the synaptonemal complex transverse filament and CG DNA hypomethylation in *zyp1 met1/+* partially suppress pericentromeric crossover reduction caused by *zyp1* or *met1/+* alone.

Next, we quantified crossovers in the sub-telomeric *CTL3.2* interval and found that, in agreement with the previous reports (Mirouze et al. 2012; Yelina et al. 2012; Yelina et al. 2015; Capilla-Pérez et al. 2021; France et al. 2021), single *met1/+* heterozygotes and *zyp1* knockouts increased sub-telomeric crossovers compared to the wild type (Wilcoxon test, *p*-value < 0.01, **Figure 1F**, **Supplemental Tables 4, 5>**). Combining *met1/+* with *zyp1* led to an additive effect with crossover frequencies further increased compared to single *met1/+* (Wilcoxon test, *p*-value = 1.9*10^−4^) or *zyp1* (Wilcoxon test, *p*-value = 3.2*10^−4^, **Figure 1F**, **Supplemental Tables 4, 5>**).

In summary, we found that *zyp1* and *met1/+* genetically interact to control sub-telomeric and pericentromeric crossovers.

### Depletion of the meiotic chromosome axis ASY1 and CG DNA hypomethylation genetically interact to control pericentromeric crossovers

To investigate whether CG DNA methylation and meiotic chromosome axis genetically interact to control crossovers, we combined *met1/+* and *asy1/+* heterozygotes and introgressed them into *CTL3.2* and *CTL3.9* via crossing. ASY1 is a HORMA domain protein, a key component of the meiotic chromosome axis (Mercier et al. 2015; Lambing et al. 2020; Pochon et al. 2023; Cai et al. 2026). We used *asy1/+* rather than *asy1* knockouts because, similarly to *asy1* null mutants, *asy1/+* exhibits pronounced crossover redistribution from pericentromeric to sub-telomeric regions; however, unlike *asy1* nulls, *asy1/+* plants remain fertile, which facilitates crossover analysis (Lambing et al. 2020).

We found that, consistent with the previous reports (Mirouze et al. 2012; Yelina et al. 2012; Yelina et al. 2015; Lambing et al. 2020), *met1/+* and *asy1/+* single heterozygotes showed reduced pericentromeric crossovers in *CTL3.9* compared to the wild type (Wilcoxon test, *p*-value < 0.01, **Figure 1G**, **Supplemental Tables 6, 7>**). *met1/+ asy1/+* double heterozygotes showed a further reduction in the pericentromeric crossovers compared to *met1/+* (Wilcoxon test *p*-value = 3.6*10^−6^) and *asy1/+* alone (Wilcoxon test *p*-value = 2.9*10^−6^, **Figure 1G**, **Supplemental Tables 6, 7>**).

Next, we quantified crossovers in the sub-telomeric *CTL3.2* interval and found that, in agreement with the previous reports (Mirouze et al. 2012; Yelina et al. 2012; Yelina et al. 2015; Lambing et al. 2020), crossovers were increased in both *met1/+* (Wilcoxon test *p*-value = 0.004, **Supplemental Tables 8, 9>**) and *asy1/+* single heterozygotes compared to the wild type (Wilcoxon test *p*-value = 0.018 **Supplemental Tables 8, 9>**), however, double *met1/+ asy1/+* heterozygotes showed very similar recombination sub-telomeric recombination frequencies compared to *met1/+* and *asy1/+* alone (Wilcoxon test, *p*-value > 0.01 **Figure 1H**, **Supplemental Tables 8, 9>**).

In summary, we found that *met1/+* and *asy1/+* act together to reduce pericentromeric crossovers, but do not show additive or synergistic effects on sub-telomeric crossovers.

### CHG DNA hypomethylation cannot compensate for the loss of pericentromeric crossovers caused by the depletion of axis or synaptonemal complex

CHG DNA hypomethylation increases recombination in Arabidopsis pericentromeres (Underwood et al. 2018), however depletion of the meiotic chromosome axis or synaptonemal complex proteins leads to a decrease in pericentromeric crossovers (Lambing et al. 2020; Capilla-Pérez et al. 2021). Here we asked whether CHG DNA hypomethylation could suppress pericentromeric crossover reduction caused by the depletion of meiotic chromosome axis or synaptonemal complex.

To test this, we combined *zyp1* with a mutation in the *CHROMOMETHYLASE3* (*CMT3*), which leads to CHG DNA hypomethylation (Chan et al. 2006) and introgressed *zyp1*, *cmt3* and *zyp1 cmt3* into *CTL3.9* and *CTL3.2*. In agreement with the previous reports (Underwood et al. 2018; Capilla-Pérez et al. 2021), pericentromeric crossovers increased in *cmt3* and decreased in *zyp1* compared to the wild type (Wilcoxon test *p*-value < 0.01, **Figure 2A**, **Supplemental Tables 10, 11>**).

**Figure 2.**
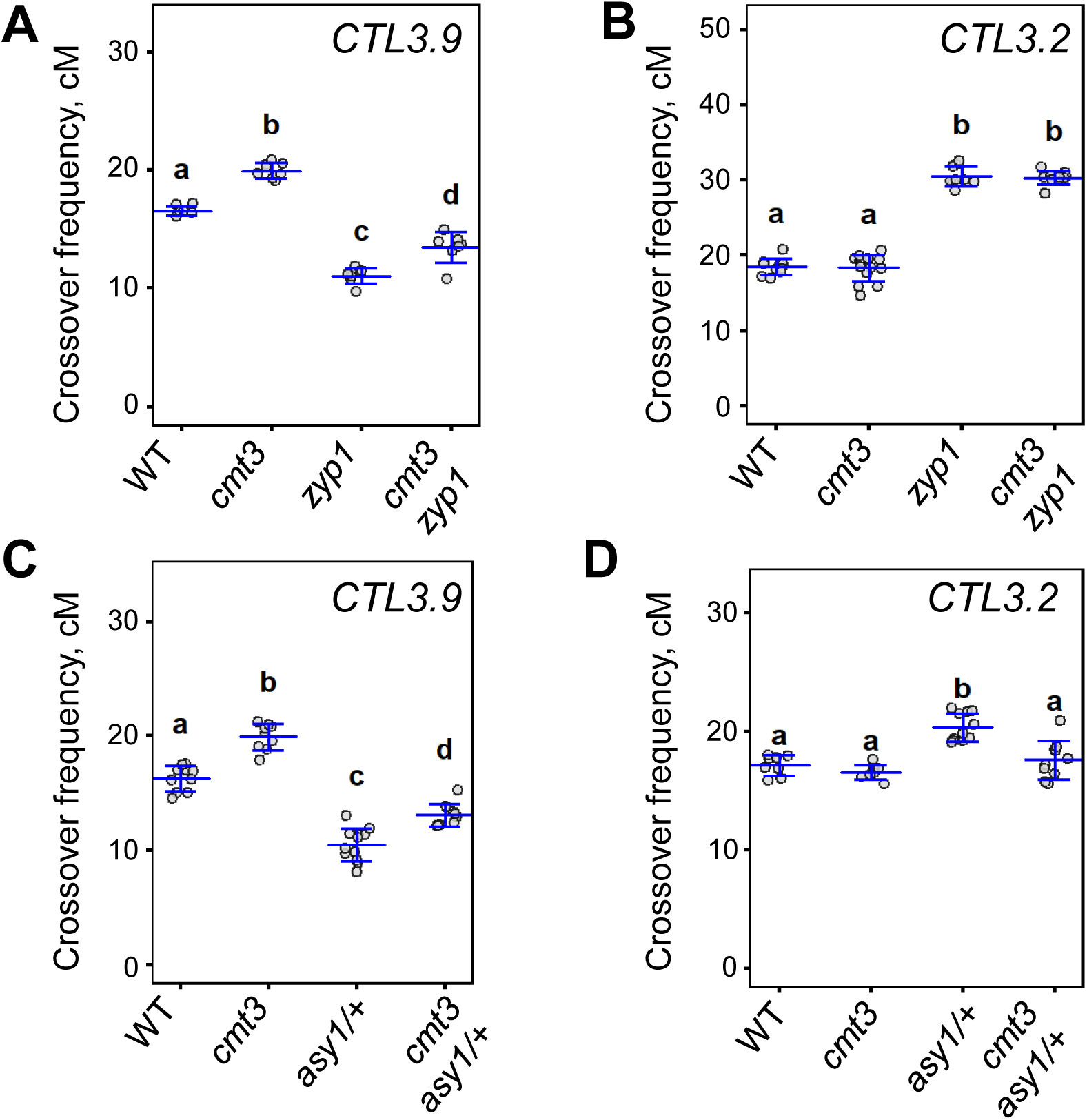
Genetic interactions between CHG DNA hypomethylation and meiotic chromosome architecture in crossover control. **(A)** Crossover frequencies (cM) in *CTL3.9* in wild type, *cmt3*, *zyp1*, *cmt3 zyp1*. Measurements from individual plants are shown as circles. Horizontal blue lines indicate means and standard deviation. Different letters indicate statistically significant differences between groups determined by a global Kruskal-Wallis test followed by pairwise Wilcoxon tests with a Holm adjustment. **(B)** As in (A) but for *CTL3.2.* **(C)** As in (A) but for wild type, *cmt3*, *asy1/+*, *cmt3 asy1/+*. **(D)** As in (C) but for *CTL3.2*.

Double *cmt3 zyp1* mutants showed reduced pericentromeric recombination compared to *cmt3* and the wild type (Wilcoxon test *p*-value = 8.8*10^−3^, **Figure 2A**, **Supplemental Tables 10, 11>**), but a modest increase compared to *zyp1* (Wilcoxon test *p*-value = 0.02, **Figure 2A**, **Supplemental Tables 10, 11>**). This suggests that CHG DNA hypomethylation is unable to fully compensate for the pericentromeric crossover reduction caused by the synaptonemal complex depletion.

We quantified crossovers in the sub-telomeric *CTL3.2* interval and found that, in agreement with the previous reports (Underwood et al. 2018; Capilla-Pérez et al. 2021; France et al. 2021), sub-telomeric crossovers were unchanged in *cmt3* (Wilcoxon test *p*-value = 1, **Figure 2B**, **Supplemental Tables 12, 13>**) and upregulated in *zyp1* compared to the wild type (Wilcoxon test *p*-value = 1.1*10^−3^, **Figure 2B**, **Supplemental Tables 12, 13>**). Double *zyp1 cmt3* mutants phenocopied *zyp1* (Wilcoxon test *p*-value = 1, **Figure 2B**, **Supplemental Tables 12, 13>**), suggesting that *zyp1* is epistatic to *cmt3* in the sub-telomeric crossover control.

Next, we combined *asy1/+* and *cmt3*, introgressed *asy1/+*, *cmt3*, and *asy1/+ cmt3* into *CTL3.9* and *CTL 3.2* and quantified crossovers in the chromosome 3 pericentromeric and sub-telomeric intervals. In agreement with the previous reports (Underwood et al. 2018; Lambing et al. 2020), *asy1/+* led to a pericentromeric crossover reduction (Wilcoxon test *p*-value = 5.2*10^−4^, **Figure 2C**, **Supplemental Tables 14, 15>**) while *cmt3* led to a pericentromeric crossover increase compared to the wild type (Wilcoxon test *p*-value = 1.1*10^−3^, **Figure 2C**, **Supplemental Tables 14, 15>**). Similarly to *zyp1 cmt3*, pericentromeric crossovers in *asy1/+ cmt3* were lower than in *cmt3* and the wild type but showed a slight but significant increase compared to *asy1/+* (Wilcoxon test *p*-value = 1.2*10^−3^, **Figure 2C**, **Supplemental Tables 14, 15>**). This suggests that CHG DNA hypomethylation is unable to fully compensate for the pericentromeric crossover reduction caused by the depletion of the chromosome axis.

In the sub-telomeric *CTL3.2* interval, in agreement with the previous reports (Underwood et al. 2018; Lambing et al. 2020) recombination frequencies between the wild type and *cmt3* showed no statistically significant differences (Wilcoxon test *p*-value = 0.3, **Figure 2D**, **Supplemental Tables 16, 17>**), while *asy1/+* showed a statistically significant crossover increase (Wilcoxon test *p*-value = 1.6*10^−3^, **Figure 2D**, **Supplemental Tables 16, 17>**). *asy1/+ cmt3* phenocopied *cmt3* (Wilcoxon test *p*-value = 0.3, **Figure 2D**, **Supplemental Tables 16, 17>**), suggesting that *cmt3* is epistatic to *asy1/+* in controlling sub-telomeric crossovers.

In summary, we found that *cmt3* modulates pericentromeric crossovers independently of *asy1/+* or *zyp1*, and CHG DNA hypomethylation is unable to fully compensate for the pericentromeric crossover reduction caused by the loss of ASY1 or ZYP1. In the sub-telomeric crossover control, *zyp1* is epistatic to *cmt3*, while *cmt3* is epistatic to *asy1/+*.

### *ASY1* overexpression increases pericentromeric crossovers

Meiotic chromosome axis and synaptonemal complex play an important role in pericentromeric crossover control. We hypothesised that if ASY1 depletion leads to a dosage-dependent decrease in pericentromeric crossovers (Lambing et al. 2020), increasing ASY1 dosage could have an opposite effect and lead to a pericentromeric crossover increase.

To test this hypothesis, we overexpressed *ASY1* under the control of its native promoter (hereinafter, *ASY1-OE*) in homozygous *CTL3.9* reporter lines. We generated fourteen independent *ASY1-OE* T_1_s, crossed them to the wild type Col-0 to bring *CTL3.9* to a hemizygous state (*CTL3.9/++*) to enable *CTL3.9* crossover measurements, selected BC_1_T_1_ progenies carrying the *ASY1-OE* transgene and measured recombination in the *CTL3.9* interval (**Figure 3A, C**, **Supplemental Table 18**). We found a statistically significant increase in crossover recombination in *CTL3.9* compared to the wild type (Wilcoxon test *p*-value = 1.15*10^−9^, **Figure 3C**, **Supplemental Tables 18, 19>**) suggesting that *ASY1* overexpression leads to an upregulation of pericentromeric crossovers. The crossover upregulation was maintained in the progenies of four independent BC_1_T_1_, carrying the *ASY1-OE* transgene, which we refer to as BC_1_F_2_ (Wilcoxon test *p*-value = 2.25*10^−5^, **Figure 3A, C**, **Supplemental Tables 18, 19>**).

**Figure 3.**
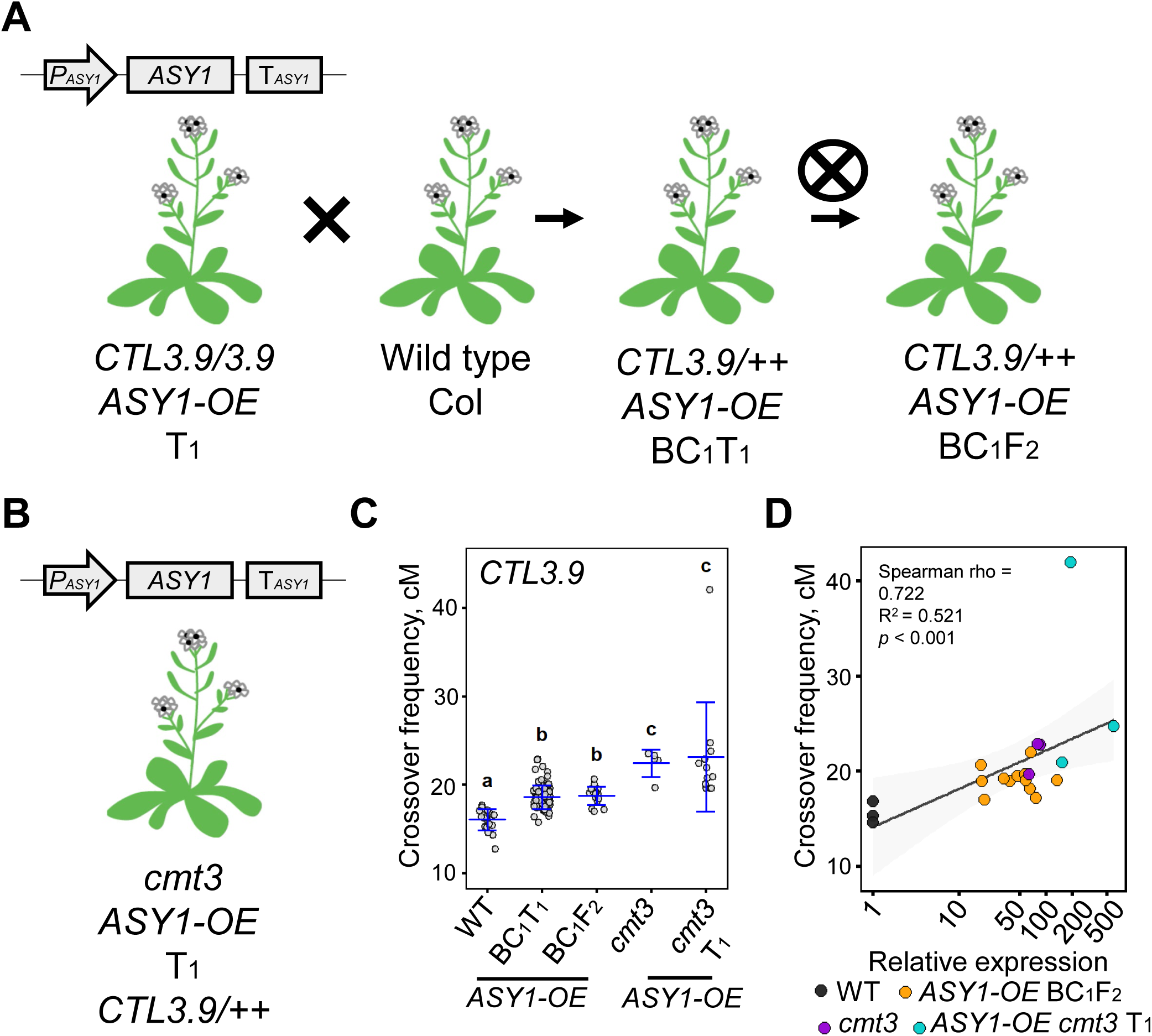
*ASY1* overexpression increases pericentromeric recombination. **(A)** A construct encoding *ASY1* under its native promoter and terminator was transformed into a homozygous *CTL3.9* reporter in Col-0 background. T_1_ transformants were selected and backcrossed to the wild type Col-0 to generate BC_1_T_1_ and enable crossover frequency quantification in the *CTL3.9* interval. Four BC_1_T_1_ were self-fertilized, three BC_1_F_2_ per each BC_1_T_1_ carrying *ASY1-OE* transgene and hemizygous *CTL3.9/++* were selected for crossover and *ASY1* transcript levels quantification. **(B)** To generate *ASY1-OE* in *cmt3* background, a construct encoding *ASY1* under its native promoter and terminator was transformed into a hemizygous *CTL3.9/++* reporter in the *cmt3* background. T_1_ transformants carrying the *ASY1-OE* transgene and hemizygous *CTL3.9/++* were selected for crossover and *ASY1* transcript levels quantification. **(C)** Crossover frequencies (cM) in *CTL3.9* in the wild type, *cmt3*, *ASY1-OE* (BC_1_T_1_, BC_1_F_2_) and *ASY1-OE* in *cmt3* (*cmt3* T_1_). Measurements from individual plants are shown as circles. Horizontal blue lines indicate means and standard deviation. Different letters indicate statistically significant differences between groups determined by a global Kruskal-Wallis test followed by pairwise Wilcoxon tests with a Holm adjustment. **(D)** Correlation plot between *CTL3.9* recombination frequencies and *ASY1* relative transcript levels. Data points represent individual samples, color-coded by genotype (wild type - charcoal; *ASY1-OE* in the wild-type (BC_1_F_2_) - amber; *cmt3* - violet; *ASY1-OE* in *cmt3* (*cmt3* T_1_) - cyan). The solid line indicates the global linear trend line with a 95% confidence interval shaded in grey. *x*-axis is displayed on a logarithmic scale. Statistical analysis was performed using Spearman’s rank-order correlation.

Next, we tested whether *ASY1* overexpression and CHG DNA hypomethylation cooperate to upregulate pericentromeric crossovers. We transformed *cmt3* carrying hemizygous *CTL3.9/++* with the *ASY1-OE* construct and selected thirteen T_1_ transformants that carried the *ASY1-OE* transgene and were hemizygous for *CTL3.9/++* to enable crossover measurements (**Figure 3B**). We found that *cmt3* showed higher *CTL3.9* recombination than the wild type and *ASY1-OE* lines in the wild type background. Twelve out of thirteen *cmt3 ASY1-OE* T1s phenocopied *cmt3* (Wilcoxon test *p*-value = 0.69, **Figure 3C**, **Supplemental Tables 18, 19>**) with only one out of the thirteen T_1_s showing a recombination frequency of 42 cM that was approximately twice as high compared to *ASY1-OE* or *cmt3* alone (**Figure 3C**, **Supplemental Tables 18, 19>**). The increase in *ASY1* transcript levels positively correlated with the increase in *CTL3.9* recombination (Spearman’s rank correlation, rho = 0.722, R^2^ = 0.521, *p*-value = 0.0002169, **Figure 3D**). We found that one out of 93 *ASY1-OE* BC_1_T_1_ that we analysed (which represent progeny of 14 independent *ASY1-OE* T_1_s in the wild type background) and one out of the thirteen *ASY1-OE* T_1_s in the *cmt3* background, showed segregation distortion of *eGFP* and *dsRed CTL3.9* reporters (**Supplemental Table 20**).

In summary, we found that increased *ASY1* dosage upregulates pericentromeric *CTL3.9* crossovers, albeit to a lower level than CHG DNA hypomethylation. We show that combining *ASY1* overexpression with CHG DNA hypomethylation does not surpass pericentromeric crossover upregulation of *cmt3*.

## Discussion

In this study, we identify additive and independent roles for meiotic chromosome architecture and cytosine DNA methylation in crossover control and demonstrate that Arabidopsis pericentromeric crossovers are limited by the dosage of a HORMA domain meiotic chromosome axis protein ASY1.

CHG DNA hypomethylation epigenetically activates Arabidopsis pericentromeric crossovers (Underwood et al. 2018), while depletion of meiotic chromosome axis and synaptonemal complex proteins ASY1 and ZYP1 downregulates recombination in Arabidopsis pericentromeres (Lambing et al. 2020; Capilla-Perez et al. 2021). We hypothesised that epigenetic crossover activation in *cmt3* could suppress pericentromeric crossover loss caused by ASY1 or ZYP1 depletion. However, this was not the case. We found that although *cmt3* retained its ability to upregulate pericentromeric crossovers when combined with *zyp1* or *asy1/+*, the epigenetic crossover activation caused by *cmt3* was insufficient to fully rescue crossover deficits imposed by *zyp1* and *asy1/+*. These data imply that although heterochromatin depletion may permit crossover formation, structural integrity of the meiotic chromosome axis is essential to drive recombination.

A reduced dosage of *ASY1* remodels Arabidopsis crossovers, shifting them from pericentromeres to sub-telomeric regions (Lambing et al. 2020). We hypothesized that increasing *ASY1* dosage would conversely upregulate pericentromeric recombination, which we confirmed experimentally. Consequently, we demonstrate a new, non-epigenetic strategy to stimulate pericentromeric crossovers in Arabidopsis by overexpressing *ASY1*.

Next, we hypothesized that combining *ASY1*-overexpression with the crossover-permissive pericentromeric chromatin of *cmt3* mutants would additively activate pericentromeric crossovers. Contrary to our expectations, this additive effect was not observed. Intriguingly, in twelve out of thirteen independent lines overexpressing *ASY1* in a CHG-hypomethylated *cmt3* background, pericentromeric recombination frequencies phenocopied the *cmt3* single mutant. This suggests that the overexpression of *ASY1* is insufficient to stimulate pericentromeric recombination beyond the threshold achieved by CHG DNA hypomethylation alone. However, the fact that one of the thirteen lines exhibited an approximate doubling of *CTL3.9* recombination compared to *cmt3* mutants and *ASY1*-overexpression lines could suggest that pericentromeric recombination capacity is not entirely saturated. Further research is needed to elucidate the regulatory factors that limit pericentromeric recombination when repressive heterochromatin is depleted and *ASY1* dosage is elevated, as well as to develop strategies to achieve this robustly.

In the future, expanding the repertoire of non-epigenetic strategies to unlock plant pericentromeres may require a multi-layered genetic approach, where in addition to overexpressing *ASY1*, dosage of other meiotic chromosome axis proteins (Ferdous et al. 2012; Chambon et al. 2018) needs to be modulated. It would be very interesting to test whether combining *ASY1* overexpression with the recently reported mutations in the cohesion establishment factor CTF18, the centromeric cohesin protector SGO2 and the deSUMOylase SPF2 (Salinas Gamboa et al. 2026) would have additive effects on upregulating pericentromeric crossovers. Another route may involve coupling altered dosage of the chromosome architecture proteins with pro-crossover *HEI10* overexpression (Ziolkowski et al. 2017), while suppressing the non-crossover pathway by mutating anti-crossover factors, including *FANCM*, *FIGL1* and *RECQ4* (Ziolkowski et al. 2017, Serra et al. 2018; Durand et al. 2022; Jing et al. 2025). A targeted approach could serve as an alternative to global modulation of meiotic gene expression to ‘unlock’ recombination in crossover-poor chromosome regions. For example, in yeast, tethering the ASY1 homolog Hop1 to crossover-suppressed regions successfully converts these coldspots into recombination hotspots (Shodhan et al. 2022). Investigating whether a similar targeted approach can be applied in plants, and comparing its efficacy to global *ASY1* overexpression, represents a promising avenue for future crossover engineering.

While *ASY1* loss can lead to genome instability and chromoanagenesis (Guo et al. 2023; Pochon et al. 2023), we observed that two out of the 27 independent *ASY1*-overexpressing lines exhibited segregation distortion of the *eGFP* and *dsRed* markers within the *CTL3.9* reporter. Further investigations are required to determine whether *ASY1* overexpression directly drives this segregation distortion and to assess whether elevated *ASY1* dosage can compromise genome stability.

While CHG hypomethylation controls the *Arabidopsis* crossover landscape independently of *zyp1* and *asy1/+*, CG hypomethylation displays additive interactions when combined with *zyp1* or *asy1/+*. This contrast is in line with the previous findings that CG and CHG DNA hypomethylation exert different effects on the Arabidopsis crossover landscapes (Mirouze et al. 2012, Yelina et al. 2012, Yelina et al. 2015; Underwood et al. 2018). We found significantly elevated pericentromeric and subtelomeric crossovers in *met1/+ zyp1* compared to single *met1/+* and *zyp1*. This additive increase is consistent with a model where *met1/+*-mediated CG hypomethylation boosts meiotic double-strand break (Choi e al. 2018) formation, but wild-type crossover interference prevents these extra breaks from maturing into crossovers (Yelina et al. 2015). By removing interference (Capilla-Pérez et al. 2021; France et al. 2021), the *zyp1* mutation licenses these additional pericentromeric DSBs to resolve as crossovers. Conversely, crossover interference remains active in *asy1/+* (Lambing et al. 2020) explaining why *met1/+asy1/+* plants do not display elevated pericentromeric recombination compared to single *met1/+* and *asy1/+*. Intriguingly, *met1/+asy1/+* double heterozygotes instead exhibit an additive reduction in pericentromeric crossovers, implying either a global suppression of crossover numbers or a compensatory remodeling shift into interstitial chromosomal intervals which were not analysed in this study.

Compared to Arabidopsis, many major crops possess substantially larger pericentromeres with a higher transposable element density (Taagen et al. 2020). These massive pericentromeric regions are predicted to harbor up to a fifth of all protein-coding genes (Taagen et al. 2020), which remain largely inaccessible to breeders due to crossover suppression. Because disrupting heterochromatin in crop species often triggers sterility and genome instability (Dorweiler et al. 2000; Parkinson et al. 2007; Gouil et al. 2016; Corem et al. 2018; Grover et al. 2018; Xu et al. 2020; Gers et al. 2026), non-epigenetic crossover activation within these ‘cold’ regions offers a highly promising alternative strategy to ‘unlock’ this untapped genetic diversity. Given the high degree of functional conservation among meiotic chromosome architecture proteins across Arabidopsis, wheat, barley, rice, and maize (Wang et al. 2021), further investigations are required to determine whether the pericentromeric crossover upregulation achieved via *ASY1* overexpression can be successfully translated into crop species. Notably, this approach may prove more feasible in diploid crops, because in polyploids, axis components such as ASY1 and ASY3 have experienced strong evolutionary selection to skew crossovers toward subtelomeric intervals, as pericentromeric crossovers in polyploid backgrounds can induce deleterious homeologous recombination and genome instability (Gonzalo et al. 2025).

In summary, our findings reveal that while permissive chromatin is necessary for recombination, meiotic chromosome architecture is the definitive gatekeeper for crossover formation. A non-epigenetic approach to enhance recombination in crossover-suppressed regions represents an important addition to the growing toolkit of strategies for overcoming pericentromeric crossover suppression, with considerable potential to optimise recombination landscapes in both model systems and major crop species.

## Data availability

*ASY1-OE* plasmid and Arabidopsis transgenic lines generated in this study are available upon request. Supplementary table 1 contains oligonucleotides used in this study, Supplementary tables 2–20 contain raw data used for crossover frequency quantification and outputs of statistical analysis.

## Acknowledgements

We thank Prof James Higgins for providing *zyp1a zyp1b* seeds and Dr Evan Ellison for providing pTRANS_210d binary vector.

---

**Supplemental Table 1.**
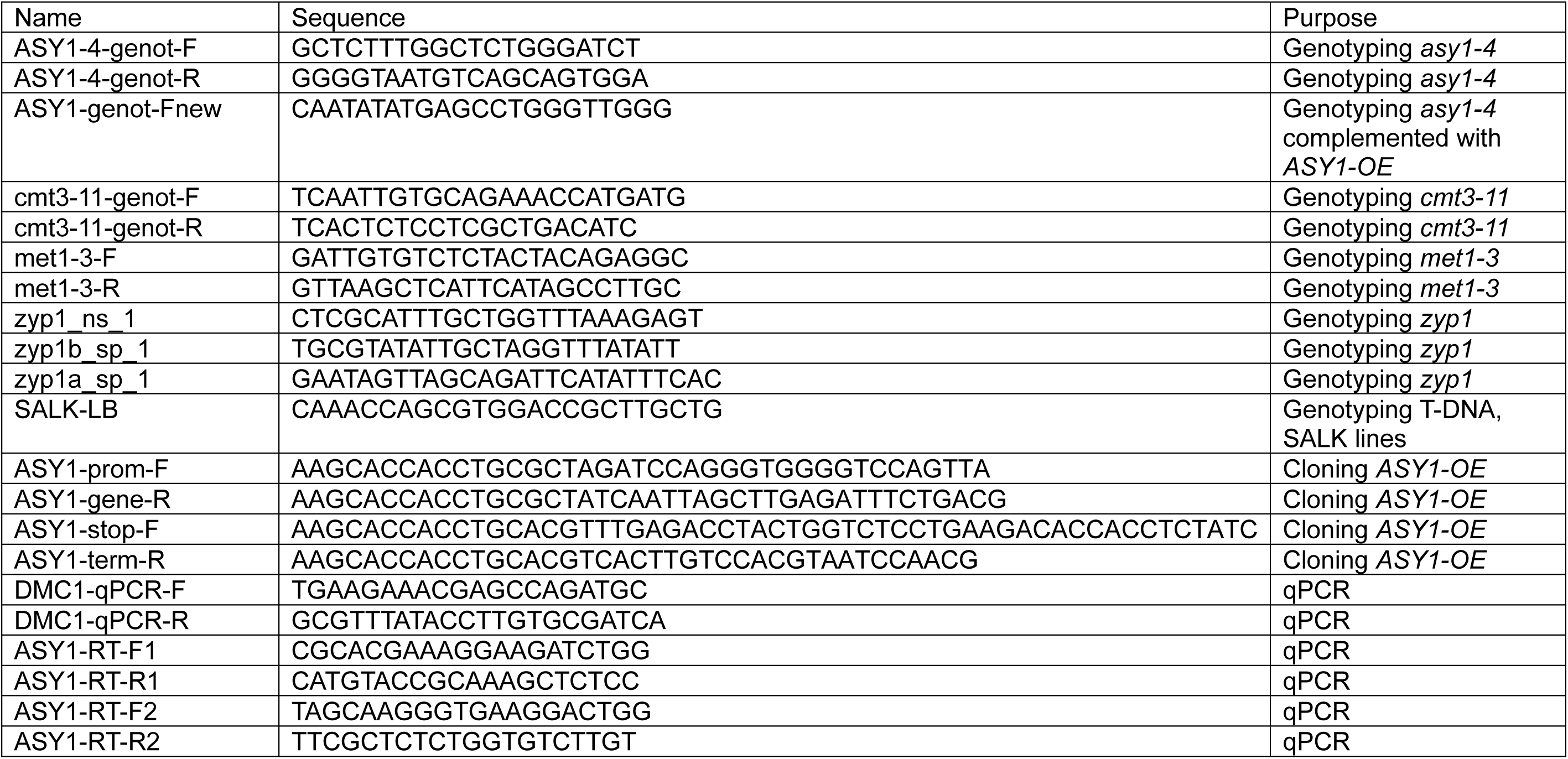
Oligonucleotides used in this study.

**Supplemental Table 2.** Genetic distances of *CTL3.9* in wild type, *zyp1*, *met1/+* and *zyp1 met1/+*. cM were calculated using the formula: cM = 100 × (1 – [1-2(*N_G_*+*N_R_*)/*N_T_*]^½^), where *N_G_* is a number of green-alone fluorescent seeds, *N_R_* is a number of red-alone fluorescent seed and *N_T_* is the total number of seeds counted.

| Genotype | Green-alone | Red-alone | Both red and green | Neither red or green | Total | *CTL3.9*, cM | Average, *CTL3.9*, cM |
| --- | --- | --- | --- | --- | --- | --- | --- |
| Wild type | 179 | 171 | 1488 | 399 | 2237 | 17.11 |  |
| Wild type | 157 | 175 | 1637 | 449 | 2418 | 14.83 |  |
| Wild type | 164 | 182 | 1577 | 429 | 2352 | 15.99 |  |
| Wild type | 146 | 187 | 1577 | 390 | 2300 | 15.71 |  |
| Wild type | 169 | 165 | 1464 | 365 | 2163 | 16.86 |  |
| Wild type | 166 | 176 | 1662 | 452 | 2456 | 15.06 |  |
| Wild type | 174 | 203 | 1604 | 371 | 2352 | 17.57 |  |
| Wild type | 161 | 126 | 1246 | 347 | 1880 | 16.65 |  |
| Wild type | 169 | 170 | 1568 | 471 | 2378 | 15.45 | 16.14 |
| *met1/+* | 109 | 147 | 1530 | 437 | 2223 | 12.27 |  |
| *met1/+* | 131 | 104 | 1577 | 475 | 2287 | 10.87 |  |
| *met1/+* | 131 | 114 | 1602 | 456 | 2303 | 11.27 |  |
| *met1/+* | 94 | 84 | 1228 | 374 | 1780 | 10.56 |  |
| *met1/+* | 70 | 54 | 855 | 252 | 1231 | 10.64 |  |
| *met1/+* | 63 | 70 | 1127 | 325 | 1585 | 8.78 |  |
| *met1/+* | 95 | 138 | 1507 | 479 | 2219 | 11.12 |  |
| *met1/+* | 94 | 110 | 1433 | 435 | 2072 | 10.38 |  |
| *met1/+* | 105 | 106 | 1357 | 395 | 1963 | 11.40 |  |
| *met1/+* | 118 | 111 | 1541 | 394 | 2164 | 11.21 |  |
| *met1/+* | 81 | 91 | 1404 | 432 | 2008 | 8.97 |  |
| *met1/+* | 134 | 106 | 1515 | 418 | 2173 | 11.73 |  |
| *met1/+* | 72 | 66 | 1051 | 318 | 1507 | 9.62 | 10.68 |
| *zyp1* | 85 | 106 | 1694 | 445 | 2330 | 8.56 |  |
| *zyp1* | 115 | 90 | 1520 | 450 | 2175 | 9.92 |  |
| *zyp1* | 81 | 105 | 1486 | 416 | 2088 | 9.34 |  |
| *zyp1* | 99 | 104 | 1486 | 434 | 2123 | 10.07 |  |
| *zyp1* | 87 | 74 | 1219 | 393 | 1773 | 9.54 |  |
| *zyp1* | 139 | 134 | 1666 | 494 | 2433 | 11.93 |  |
| *zyp1* | 97 | 115 | 1337 | 349 | 1898 | 11.87 |  |
| *zyp1* | 129 | 118 | 1521 | 459 | 2227 | 11.79 | 10.38 |
| *zyp1 met1/+* | 135 | 142 | 1510 | 395 | 2182 | 13.62 |  |
| *zyp1 met1/+* | 129 | 132 | 1384 | 360 | 2005 | 14.00 |  |
| *zyp1 met1/+* | 137 | 152 | 1544 | 388 | 2221 | 13.99 |  |
| *zyp1 met1/+* | 90 | 83 | 1005 | 297 | 1475 | 12.51 |  |
| *zyp1 met1/+* | 144 | 125 | 1559 | 424 | 2252 | 12.76 |  |
| *zyp1 met1/+* | 144 | 130 | 1499 | 434 | 2207 | 13.30 |  |
| *zyp1 met1/+* | 136 | 96 | 1366 | 368 | 1966 | 12.59 |  |
| *zyp1 met1/+* | 144 | 140 | 1519 | 423 | 2226 | 13.70 |  |
| *zyp1 met1/+* | 118 | 153 | 1477 | 434 | 2182 | 13.30 |  |
| *zyp1 met1/+* | 143 | 131 | 1554 | 424 | 2252 | 13.01 |  |
| *zyp1 met1/+* | 128 | 116 | 1552 | 440 | 2236 | 11.58 |  |
| *zyp1 met1/+* | 106 | 136 | 1423 | 423 | 2088 | 12.35 |  |
| *zyp1 met1/+* | 103 | 95 | 1127 | 340 | 1665 | 12.70 | 13.03 |

**Supplemental Table 3.** Statistical significance (Mann-Whitney test) of genetic distances of *CTL3.9* in wild type, *zyp1*, *met1/+* and *zyp1 met1/+*.

|  | Wild type | *met1/+* | *zyp1* |
| --- | --- | --- | --- |
| *met1/+* | 0.0005348 | n/a | n/a |
| *zyp1* | 0.0013265 | 0.69040 | n/a |
| *zyp1 met1/+* | 0.0005348 | 0.000156 | 0.001327 |

**Supplemental Table 4.** Genetic distances of *CTL3.2* in wild type, *zyp1*, *met1/+* and *zyp1 met1/+*. cM were calculated using the formula: cM = 100 × (1 – [1-2(*N_G_*+*N_R_*)/*N_T_*]^½^), where *N_G_* is a number of green-alone fluorescent seeds, *N_R_* is a number of red-alone fluorescent seed and *N_T_* is the total number of seeds counted.

| Genotype | Green-alone | Red-alone | Both red and green | Neither red or green | Total | CTL3.2, cM | Average, CTL3.2, cM |
| --- | --- | --- | --- | --- | --- | --- | --- |
| Wild type | 175 | 186 | 1690 | 402 | 2453 | 16 |  |
| Wild type | 157 | 146 | 1416 | 356 | 2075 | 15.86 |  |
| Wild type | 192 | 181 | 1696 | 438 | 2507 | 16.19 |  |
| Wild type | 187 | 198 | 1374 | 346 | 2105 | 20.36 |  |
| Wild type | 212 | 173 | 1570 | 393 | 2348 | 18.02 |  |
| Wild type | 176 | 198 | 1585 | 392 | 2351 | 17.43 |  |
| Wild type | 208 | 210 | 1523 | 395 | 2336 | 19.87 |  |
| Wild type | 201 | 207 | 1643 | 375 | 2426 | 18.54 |  |
| Wild type | 159 | 144 | 1263 | 336 | 1902 | 17.45 |  |
| Wild type | 206 | 200 | 1753 | 413 | 2572 | 17.28 |  |
| Wild type | 178 | 185 | 1451 | 416 | 2230 | 17.88 |  |
| Wild type | 193 | 183 | 1509 | 425 | 2310 | 17.87 | 17.73 |
| *met1/+* | 236 | 194 | 1328 | 326 | 2084 | 23.36 |  |
| *met1/+* | 222 | 207 | 1227 | 323 | 1979 | 24.74 |  |
| *met1/+* | 209 | 183 | 1166 | 292 | 1850 | 24.09 |  |
| *met1/+* | 283 | 241 | 1452 | 332 | 2308 | 26.11 |  |
| *met1/+* | 290 | 255 | 1487 | 339 | 2371 | 26.5 |  |
| *met1/+* | 285 | 244 | 1545 | 337 | 2411 | 25.09 |  |
| *met1/+* | 245 | 212 | 1303 | 290 | 2050 | 25.56 |  |
| *met1/+* | 190 | 184 | 1121 | 306 | 1801 | 23.54 |  |
| *met1/+* | 97 | 117 | 757 | 186 | 1157 | 20.62 |  |
| *met1/+* | 179 | 207 | 1406 | 323 | 2115 | 20.31 |  |
| *met1/+* | 225 | 266 | 1594 | 410 | 2495 | 22.13 | 23.82 |
| *zyp1* | 297 | 287 | 1383 | 232 | 2199 | 31.53 |  |
| *zyp1* | 256 | 225 | 1139 | 300 | 1920 | 29.36 |  |
| *zyp1* | 293 | 264 | 1437 | 275 | 2269 | 28.65 |  |
| *zyp1* | 263 | 300 | 1401 | 305 | 2269 | 29.02 |  |
| *zyp1* | 403 | 206 | 1367 | 278 | 2254 | 32.2 |  |
| *zyp1* | 299 | 316 | 1436 | 269 | 2320 | 31.46 |  |
| *zyp1* | 285 | 264 | 1290 | 232 | 2071 | 31.46 |  |
| *zyp1* | 307 | 284 | 1534 | 275 | 2400 | 28.76 |  |
| *zyp1* | 319 | 302 | 1434 | 340 | 2395 | 30.62 | 30.34 |
| *zyp1 met1/+* | 277 | 340 | 1343 | 268 | 2228 | 33.21 |  |
| *zyp1 met1/+* | 283 | 294 | 1243 | 237 | 2057 | 33.74 |  |
| *zyp1 met1/+* | 268 | 232 | 1075 | 211 | 1786 | 33.66 |  |
| *zyp1 met1/+* | 352 | 247 | 1246 | 313 | 2158 | 33.30 |  |
| *zyp1 met1/+* | 310 | 221 | 988 | 191 | 1710 | 38.44 |  |
| *zyp1 met1/+* | 291 | 303 | 1278 | 215 | 2087 | 34.37 |  |
| *zyp1 met1/+* | 271 | 323 | 1266 | 329 | 2189 | 32.38 |  |
| *zyp1 met1/+* | 368 | 341 | 1436 | 253 | 2398 | 36.07 |  |
| *zyp1 met1/+* | 244 | 267 | 1143 | 222 | 1876 | 32.53 |  |
| *zyp1 met1/+* | 305 | 337 | 1401 | 225 | 2268 | 34.13 |  |
| *zyp1 met1/+* | 270 | 329 | 1244 | 243 | 2086 | 34.75 |  |
| *zyp1 met1/+* | 214 | 282 | 1003 | 203 | 1702 | 35.41 |  |
| *zyp1 met1/+* | 312 | 321 | 1382 | 279 | 2294 | 33.06 | 34.24 |

**Supplemental Table 5.** Statistical significance (Mann-Whitney test) of genetic distances of *CTL3.2* in wild type, *zyp1*, *met1/+* and *zyp1 met1/+*.

|  | **Wild type** | ***met1/+*** | ***zyp1*** |
| --- | --- | --- | --- |
| ***met1/+*** | 0.000288 | n/a | n/a |
| ***zyp1*** | 0.000321 | 0.000321 | n/a |
| ***zyp1 met1/+*** | 0.000150 | 0.000195 | 0.000321 |

**Supplemental Table 6.** Genetic distances of *CTL3.9* in wild type, *asy1/+*, *met1/+* and *asy1/+ met1/+*. cM were calculated using the formula: cM = 100 × (1 – [1-2(*N_G_*+*N_R_*)/*N_T_*] ^½^), where *N_G_* is a number of green-alone fluorescent seeds, *N_R_* is a number of red-alone fluorescent seed and *N_T_* is the total number of seeds counted.

| **Genotype** | **Green-alone** | **Red-alone** | **Both red and green** | **Neither red or green** | **Total** | ***CTL3.9*, cM** | **Average *CTL3.9*, cM** |
| --- | --- | --- | --- | --- | --- | --- | --- |
| Wild type | 162 | 187 | 1472 | 405 | 2226 | 17.15 |  |
| Wild type | 157 | 182 | 1429 | 389 | 2157 | 17.19 |  |
| Wild type | 169 | 194 | 1463 | 396 | 2222 | 17.95 |  |
| Wild type | 159 | 168 | 1524 | 418 | 2269 | 15.63 |  |
| Wild type | 188 | 140 | 1319 | 358 | 2005 | 17.97 |  |
| Wild type | 135 | 249 | 1530 | 442 | 2356 | 17.90 |  |
| Wild type | 204 | 141 | 1440 | 385 | 2170 | 17.42 |  |
| Wild type | 134 | 142 | 1368 | 374 | 2018 | 14.77 |  |
| Wild type | 207 | 170 | 1378 | 354 | 2109 | 19.84 | 17.31 |
| *met1/+* | 33 | 32 | 596 | 179 | 840 | 8.06 |  |
| *met1/+* | 66 | 95 | 1444 | 357 | 1962 | 8.57 |  |
| *met1/+* | 100 | 75 | 1350 | 394 | 1919 | 9.58 |  |
| *met1/+* | 102 | 99 | 1704 | 485 | 2390 | 8.80 |  |
| *met1/+* | 85 | 67 | 1282 | 385 | 1819 | 8.74 |  |
| *met1/+* | 96 | 85 | 1404 | 429 | 2014 | 9.43 |  |
| *met1/+* | 114 | 79 | 1519 | 473 | 2185 | 9.26 |  |
| *met1/+* | 84 | 81 | 1249 | 365 | 1779 | 9.75 |  |
| *met1/+* | 93 | 90 | 1567 | 490 | 2240 | 8.53 |  |
| *met1/+* | 79 | 89 | 1448 | 422 | 2038 | 8.61 |  |
| *met1/+* | 90 | 61 | 1401 | 406 | 1958 | 8.03 |  |
| *met1/+* | 99 | 183 | 1760 | 498 | 2540 | 11.80 |  |
| *met1/+* | 101 | 105 | 1342 | 399 | 1947 | 11.21 |  |
| *met1/+* | 144 | 99 | 1656 | 494 | 2393 | 10.73 |  |
| *met1/+* | 125 | 76 | 1430 | 444 | 2075 | 10.21 |  |
| *met1/+* | 130 | 134 | 1773 | 513 | 2550 | 10.95 | 9.52 |
| *asy1/+* | 119 | 105 | 1769 | 539 | 2532 | 9.28 |  |
| *asy1/+* | 113 | 111 | 1774 | 525 | 2523 | 9.31 |  |
| *asy1/+* | 92 | 129 | 1670 | 492 | 2383 | 9.75 |  |
| *asy1/+* | 103 | 124 | 1661 | 496 | 2384 | 10.02 |  |
| *asy1/+* | 104 | 85 | 1643 | 508 | 2340 | 8.43 |  |
| *asy1/+* | 103 | 119 | 1669 | 527 | 2418 | 9.65 |  |
| *asy1/+* | 108 | 123 | 1710 | 527 | 2468 | 9.84 |  |
| *asy1/+* | 117 | 108 | 1646 | 491 | 2362 | 10.03 |  |
| *asy1/+* | 175 | 167 | 1633 | 431 | 2406 | 15.40 |  |
| *asy1/+* | 107 | 125 | 1572 | 522 | 2326 | 10.53 |  |
| *asy1/+* | 121 | 120 | 1740 | 492 | 2473 | 10.27 |  |
| *asy1/+* | 88 | 92 | 1294 | 429 | 1903 | 9.95 |  |
| *asy1/+* | 83 | 116 | 1494 | 475 | 2168 | 9.64 |  |
| *asy1/+* | 100 | 105 | 1651 | 444 | 2300 | 9.35 |  |
| *asy1/+* | 100 | 113 | 1728 | 512 | 2453 | 9.10 |  |
| *asy1/+* | 104 | 110 | 1729 | 471 | 2414 | 9.30 |  |
| *asy1/+* | 107 | 122 | 1729 | 493 | 2451 | 9.83 | 9.98 |
| *asy1/+ met1/+* | 66 | 62 | 1599 | 475 | 2202 | 5.99 |  |
| *asy1/+ met1/+* | 42 | 61 | 1247 | 393 | 1743 | 6.10 |  |
| *asy1/+ met1/+* | 21 | 29 | 572 | 194 | 816 | 6.33 |  |
| *asy1/+ met1/+* | 43 | 57 | 1274 | 402 | 1776 | 5.80 |  |
| *asy1/+ met1/+* | 68 | 41 | 1584 | 509 | 2202 | 5.08 |  |
| *asy1/+ met1/+* | 55 | 60 | 1543 | 462 | 2120 | 5.58 |  |
| *asy1/+ met1/+* | 58 | 55 | 1300 | 388 | 1801 | 6.48 |  |
| *asy1/+ met1/+* | 74 | 48 | 1523 | 478 | 2123 | 5.92 |  |
| *asy1/+ met1/+* | 59 | 75 | 1607 | 512 | 2253 | 6.14 |  |
| *asy1/+ met1/+* | 60 | 56 | 1417 | 476 | 2009 | 5.95 |  |
| *asy1/+ met1/+* | 37 | 28 | 784 | 247 | 1096 | 6.12 |  |
| *asy1/+ met1/+* | 70 | 66 | 1579 | 470 | 2185 | 6.43 |  |
| *asy1/+ met1/+* | 80 | 72 | 1603 | 476 | 2231 | 7.06 |  |
| *asy1/+ met1/+* | 63 | 56 | 1597 | 501 | 2217 | 5.52 |  |
| *asy1/+ met1/+* | 93 | 58 | 1629 | 510 | 2290 | 6.83 |  |
| *asy1/+ met1/+* | 115 | 60 | 1733 | 556 | 2464 | 7.37 |  |
| *asy1/+ met1/+* | 77 | 63 | 1543 | 464 | 2147 | 6.75 |  |
| *asy1/+ met1/+* | 87 | 74 | 1868 | 560 | 2589 | 6.43 | 6.22 |

**Supplemental Table 7.** Statistical significance (Mann-Whitney test) of genetic distances of *CTL3.9* in wild type, *asy1/+*, *met1/+* and *asy1/+ met1/+*.

|  | **Wild type** | ***met1/+*** | ***asy1/+*** |
| --- | --- | --- | --- |
| ***met1/+*** | 0.000155 | n/a | n/a |
| ***asy1/+*** | 0.000155 | 0.2798 | n/a |
| ***asy1/+ met1/+*** | 0.000138 | $3.68 \times 10^{-6}$ | $2.89 \times 10^{-6}$ |

**Supplemental Table 8.** Genetic distances of *CTL3.2* in wild type, *asy1/+*, *met1/+* and *asy1/+ met1/+*. cM were calculated using the formula: cM = 100 × (1 – [1-2(*N_G_*+*N_R_*)/*N_T_*] ^½^), where *N_G_* is a number of green-alone fluorescent seeds, *N_R_* is a number of red-alone fluorescent seed and *N_T_* is the total number of seeds counted.

| Genotype | Green-alone | Red-alone | Both red and green | Neither red or green | Total | CTL3.2, cM | Average CTL3.2, cM |
| --- | --- | --- | --- | --- | --- | --- | --- |
| Wild type | 205 | 182 | 1478 | 344 | 2209 | 19.40 |  |
| Wild type | 193 | 184 | 1509 | 424 | 2310 | 17.93 |  |
| Wild type | 204 | 186 | 1437 | 335 | 2162 | 20.05 |  |
| Wild type | 189 | 174 | 1220 | 300 | 1883 | 21.61 |  |
| Wild type | 202 | 205 | 1637 | 382 | 2426 | 18.49 |  |
| Wild type | 174 | 191 | 1454 | 411 | 2230 | 17.99 |  |
| Wild type | 208 | 172 | 1386 | 339 | 2105 | 20.07 | 19.36 |
| *met1/+* | 203 | 262 | 1577 | 321 | 2363 | 22.13 |  |
| *met1/+* | 265 | 204 | 1459 | 336 | 2264 | 23.47 |  |
| *met1/+* | 91 | 110 | 712 | 147 | 1060 | 21.21 |  |
| *met1/+* | 200 | 206 | 1420 | 333 | 2159 | 21.01 |  |
| *met1/+* | 213 | 193 | 1426 | 323 | 2155 | 21.06 |  |
| *met1/+* | 163 | 145 | 962 | 233 | 1503 | 23.18 |  |
| *met1/+* | 255 | 186 | 1320 | 323 | 2084 | 24.05 | 22.30 |
| *asy1/+* | 207 | 170 | 1378 | 354 | 2109 | 19.84 |  |
| *asy1/+* | 189 | 174 | 1220 | 300 | 1883 | 21.61 |  |
| *asy1/+* | 179 | 217 | 1332 | 337 | 2065 | 21.48 |  |
| *asy1/+* | 173 | 216 | 1389 | 343 | 2121 | 20.43 |  |
| *asy1/+* | 186 | 242 | 1408 | 307 | 2143 | 22.50 |  |
| *asy1/+* | 275 | 250 | 1455 | 364 | 2344 | 25.70 |  |
| *asy1/+* | 258 | 223 | 1536 | 345 | 2362 | 23.01 | 22.08 |
| *asy1/+ met1/+* | 222 | 227 | 1467 | 348 | 2264 | 22.32 |  |
| *asy1/+ met1/+* | 191 | 211 | 1415 | 315 | 2132 | 21.08 |  |
| *asy1/+ met1/+* | 194 | 191 | 1425 | 322 | 2132 | 20.07 |  |
| *asy1/+ met1/+* | 232 | 192 | 1265 | 290 | 1979 | 24.40 |  |
| *asy1/+ met1/+* | 85 | 102 | 691 | 150 | 1028 | 20.24 |  |
| *asy1/+ met1/+* | 267 | 182 | 1668 | 401 | 2518 | 19.79 |  |
| *asy1/+ met1/+* | 199 | 239 | 1649 | 346 | 2433 | 20.00 | 21.13 |

**Supplemental Table 9.** Statistical significance (Mann-Whitney test) of genetic distances of *CTL3.2* in wild type, *asy1/+*, *met1/+* and *asy1/+ met1/+*.

|  | **Wild type** | ***met1/+*** | ***asy1/+*** |
| --- | --- | --- | --- |
| ***met1/+*** | 0.004 | n/a | n/a |
| ***asy1/+*** | 0.018 | 0.70148 | n/a |
| ***asy1/+ met1/+*** | 0.064 | 0.165 | 0.259 |

**Supplemental Table 10.** Genetic distances of *CTL3.9* in wild type, *cmt3*, *zyp1* and *cmt3 zyp1*. cM were calculated using the formula: cM = 100 × (1 – [1-2(*N_G_*+*N_R_*)/*N_T_*]^½^), where *N_G_* is a number of green-alone fluorescent seeds, *N_R_* is a number of red-alone fluorescent seed and *N_T_* is the total number of seeds counted.

| Genotype | Green-alone | Red-alone | Both red and green | Neither red or green | Total | CTL3.9, cM | Average, CTL3.9, cM |
| --- | --- | --- | --- | --- | --- | --- | --- |
| Wild type | 153 | 203 | 1557 | 363 | 2276 | 17.1 |  |
| Wild type | 140 | 142 | 1279 | 309 | 1870 | 16.43 |  |
| Wild type | 156 | 153 | 1315 | 348 | 1972 | 17.14 |  |
| Wild type | 144 | 108 | 1149 | 302 | 1703 | 16.09 |  |
| Wild type | 184 | 159 | 1564 | 362 | 2269 | 16.47 |  |
| Wild type | 200 | 195 | 1814 | 413 | 2622 | 16.41 |  |
| Wild type | 160 | 171 | 1458 | 394 | 2183 | 16.53 | 16.60 |
| *cmt3* | 191 | 197 | 1368 | 358 | 2114 | 20.44 |  |
| *cmt3* | 240 | 214 | 1528 | 445 | 2427 | 20.89 |  |
| *cmt3* | 209 | 222 | 1594 | 410 | 2435 | 19.63 |  |
| *cmt3* | 218 | 202 | 1571 | 444 | 2435 | 19.07 |  |
| *cmt3* | 214 | 235 | 1551 | 469 | 2469 | 20.23 |  |
| *cmt3* | 151 | 171 | 1183 | 310 | 1815 | 19.68 |  |
| *cmt3* | 152 | 246 | 1475 | 288 | 2161 | 20.52 |  |
| *cmt3* | 169 | 210 | 1432 | 361 | 2172 | 19.31 | 19.97 |
| *zyp1* | 116 | 113 | 1429 | 394 | 2052 | 11.86 |  |
| *zyp1* | 99 | 116 | 1447 | 435 | 2097 | 10.84 |  |
| *zyp1* | 133 | 137 | 1742 | 503 | 2515 | 11.38 |  |
| *zyp1* | 134 | 144 | 1880 | 517 | 2675 | 11 |  |
| *zyp1* | 98 | 95 | 1213 | 378 | 1784 | 11.48 |  |
| *zyp1* | 84 | 130 | 1657 | 443 | 2314 | 9.72 |  |
| *zyp1* | 110 | 143 | 1711 | 426 | 2390 | 11.21 | 11.07 |
| *cmt3 zyp1* | 152 | 130 | 1546 | 381 | 2209 | 13.71 |  |
| *cmt3 zyp1* | 134 | 130 | 1403 | 347 | 2014 | 14.1 |  |
| *cmt3 zyp1* | 178 | 155 | 1652 | 423 | 2408 | 14.95 |  |
| *cmt3 zyp1* | 96 | 176 | 1488 | 334 | 2094 | 13.96 |  |
| *cmt3 zyp1* | 108 | 103 | 1187 | 319 | 1717 | 13.15 |  |
| *cmt3 zyp1* | 135 | 80 | 1476 | 411 | 2102 | 10.81 |  |
| *cmt3 zyp1* | 144 | 141 | 1526 | 437 | 2248 | 13.6 | 13.47 |

**Supplemental Table 11.** Statistical significance (Mann-Whitney test) of genetic distances of *CTL3.9* in wild type, *cmt3*, *zyp1* and *cmt3 zyp1*.

|  | **Wild type** | ***cmt3*** | ***zyp1*** |
| --- | --- | --- | --- |
| ***cmt3*** | 0.00876 | n/a | n/a |
| ***zyp1*** | 0.00876 | 0.00876 | n/a |
| ***zyp1 cmt3*** | 0.00876 | 0.00876 | 0.02145 |

**Supplemental Table 12.** Genetic distances of *CTL3.2* in wild type, *cmt3*, *zyp1* and *cmt3 zyp1*. cM were calculated using the formula: cM = 100 × (1 – [1-2(*N_G_*+*N_R_*)/*N_T_*]^½^), where *N_G_* is a number of green-alone fluorescent seeds, *N_R_* is a number of red-alone fluorescent seed and *N_T_* is the total number of seeds counted.

| Genotype | Green-alone | Red-alone | Both red and green | Neither red or green | Total | *CTL3.2*, cM | Average, *CTL3.2*, cM |
| --- | --- | --- | --- | --- | --- | --- | --- |
| Wild type | 202 | 183 | 1498 | 369 | 2252 | 18.88 |  |
| Wild type | 215 | 180 | 1539 | 386 | 2320 | 18.79 |  |
| Wild type | 200 | 182 | 1471 | 359 | 2212 | 19.09 |  |
| Wild type | 206 | 186 | 1526 | 373 | 2291 | 18.9 |  |
| Wild type | 179 | 199 | 1542 | 367 | 2287 | 18.18 |  |
| Wild type | 211 | 199 | 1450 | 345 | 2205 | 20.75 |  |
| Wild type | 198 | 185 | 1570 | 354 | 2307 | 18.27 |  |
| Wild type | 184 | 177 | 1595 | 348 | 2304 | 17.14 |  |
| Wild type | 183 | 175 | 1600 | 346 | 2304 | 16.98 |  |
| Wild type | 184 | 189 | 1580 | 351 | 2304 | 17.77 | 18.47 |
| *cmt3* | 159 | 158 | 1614 | 409 | 2340 | 14.61 |  |
| *cmt3* | 199 | 177 | 1325 | 333 | 2034 | 20.61 |  |
| *cmt3* | 201 | 232 | 1632 | 385 | 2450 | 19.59 |  |
| *cmt3* | 157 | 180 | 1341 | 351 | 2029 | 18.28 |  |
| *cmt3* | 179 | 187 | 1457 | 376 | 2199 | 18.32 |  |
| *cmt3* | 175 | 186 | 1627 | 479 | 2467 | 15.9 |  |
| *cmt3* | 205 | 162 | 1373 | 354 | 2094 | 19.41 |  |
| *cmt3* | 154 | 187 | 1321 | 306 | 1968 | 19.16 |  |
| *cmt3* | 170 | 143 | 1432 | 393 | 2138 | 15.9 |  |
| *cmt3* | 171 | 192 | 1532 | 363 | 2258 | 17.63 |  |
| *cmt3* | 179 | 154 | 1271 | 316 | 1920 | 19.18 | 18.33 |
| *zyp1* | 288 | 302 | 1456 | 278 | 2324 | 29.84 |  |
| *zyp1* | 283 | 295 | 1328 | 243 | 2149 | 32.02 |  |
| *zyp1* | 332 | 325 | 1578 | 344 | 2579 | 29.96 |  |
| *zyp1* | 338 | 293 | 1428 | 297 | 2356 | 31.86 |  |
| *zyp1* | 289 | 312 | 1465 | 288 | 2354 | 30.04 |  |
| *zyp1* | 313 | 331 | 1437 | 280 | 2361 | 32.59 |  |
| *zyp1* | 313 | 270 | 1421 | 282 | 2286 | 30 |  |
| *zyp1* | 261 | 289 | 1387 | 307 | 2244 | 28.6 |  |
| *zyp1* | 294 | 276 | 1382 | 293 | 2245 | 29.84 | 30.53 |
| *cmt3 zyp1* | 308 | 290 | 1415 | 273 | 2286 | 30.95 |  |
| *cmt3 zyp1* | 275 | 317 | 1415 | 291 | 2298 | 30.37 |  |
| *cmt3 zyp1* | 274 | 295 | 1376 | 258 | 2203 | 30.47 |  |
| *cmt3 zyp1* | 271 | 319 | 1405 | 284 | 2279 | 30.56 |  |
| *cmt3 zyp1* | 260 | 230 | 1271 | 258 | 2019 | 28.26 |  |
| *cmt3 zyp1* | 252 | 242 | 1175 | 256 | 1925 | 30.23 |  |
| *cmt3 zyp1* | 278 | 305 | 1337 | 265 | 2185 | 31.71 |  |
| *cmt3 zyp1* | 304 | 262 | 1392 | 241 | 2199 | 30.34 |  |
| *cmt3 zyp1* | 254 | 253 | 1238 | 243 | 1988 | 30 | 30.32 |

**Supplemental Table 13.**
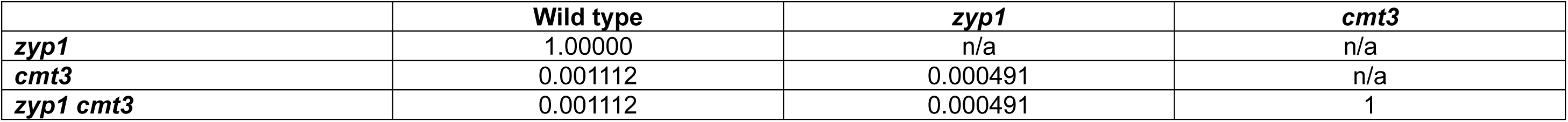
Statistical significance (Mann-Whitney test) of genetic distances of *CTL3.2* in wild type, *cmt3*, *zyp1* and *cmt3 zyp1*.

|  | **Wild type** | ***zyp1*** | ***cmt3*** |
| --- | --- | --- | --- |
| ***zyp1*** | 1.00000 | n/a | n/a |
| ***cmt3*** | 0.001112 | 0.000491 | n/a |
| ***zyp1 cmt3*** | 0.001112 | 0.000491 | 1 |

**Supplemental Table 14.**
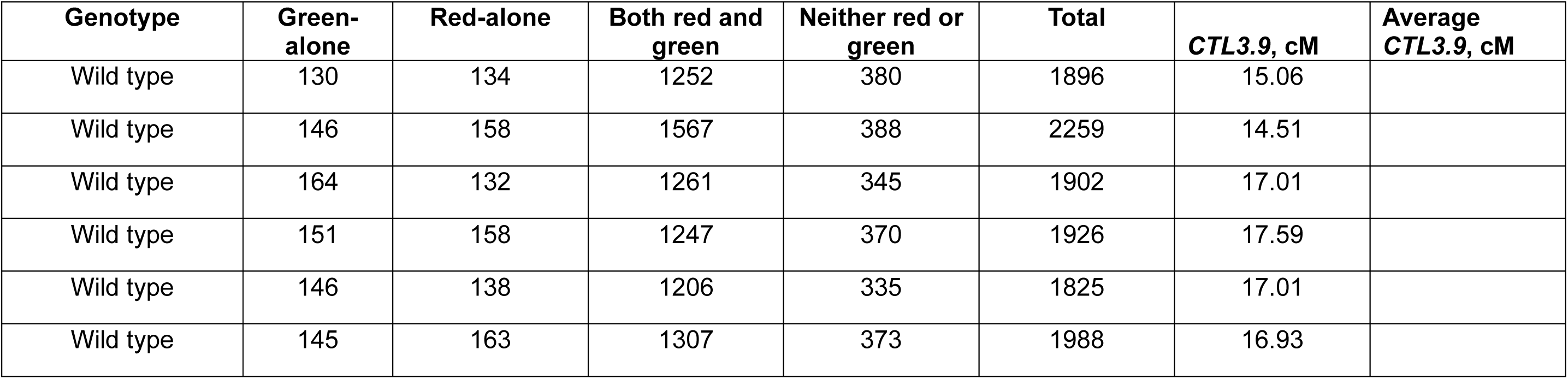
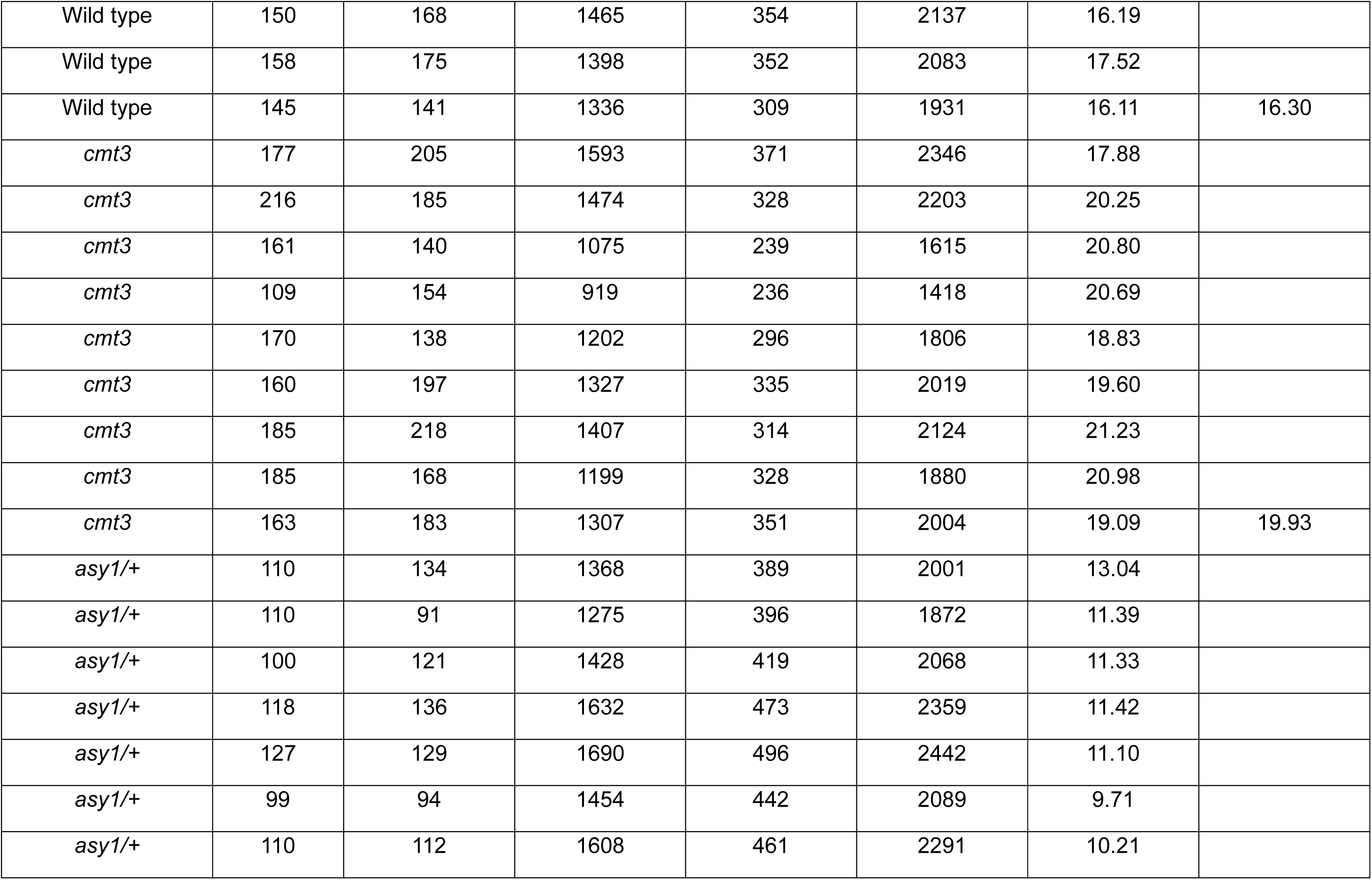
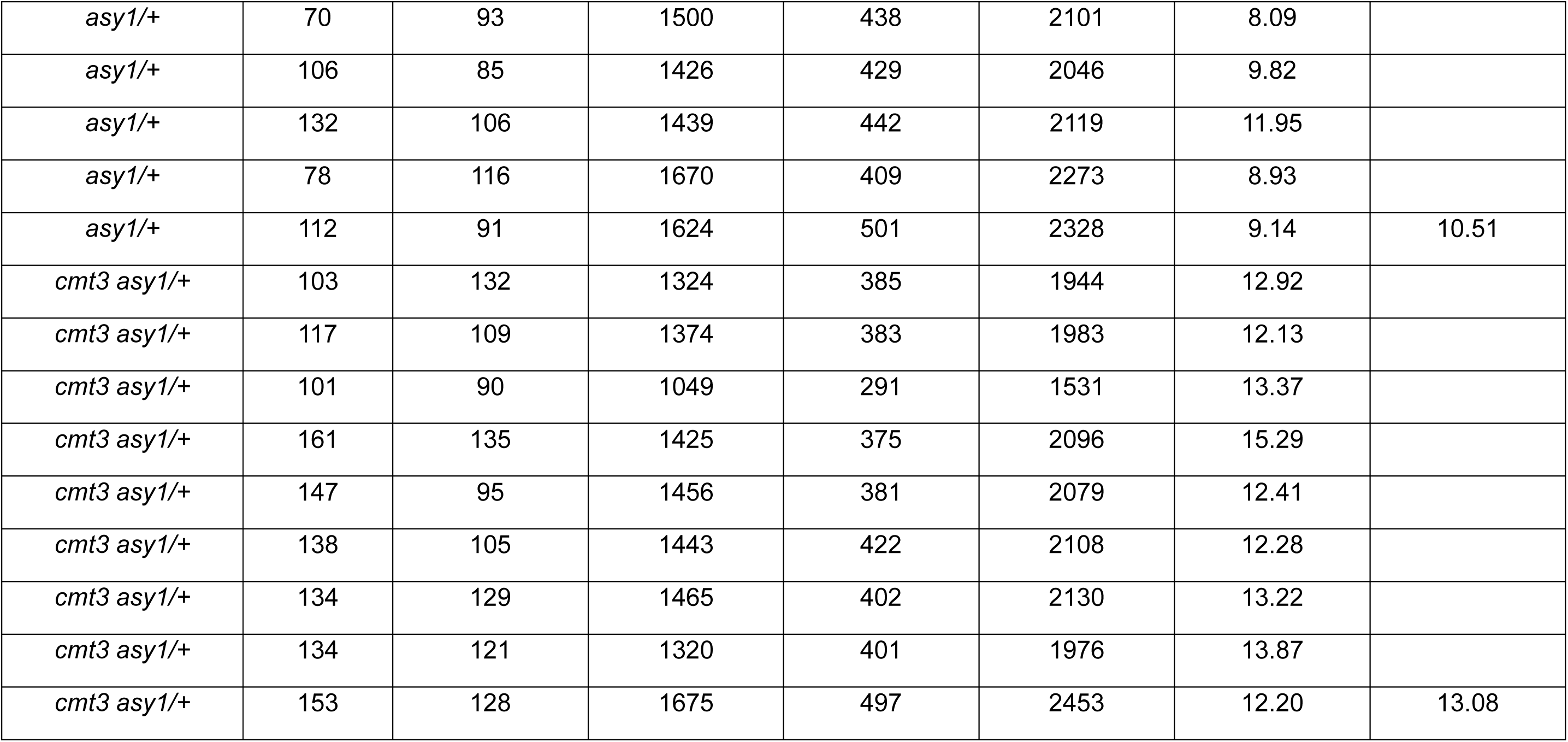
Genetic distances of *CTL3.9* in wild type, *cmt3*, *asy1/+* and *cmt3 asy1/+*. cM were calculated using the formula: cM = 100 × (1 – [1-2(*N_G_*+*N_R_*)/*N_T_*]^½^), where *N_G_* is a number of green-alone fluorescent seeds, *N_R_* is a number of red-alone fluorescent seed and *N_T_* is the total number of seeds counted.

**Supplemental Table 15.** Statistical significance (Mann-Whitney test) of genetic distances of *CTL3.9* in wild type, *cmt3*, *asy1/+* and *cmt3 asy1/+*.

|  | **Wild type** | ***cmt3*** | ***asy1/+*** |
| --- | --- | --- | --- |
| ***cmt3*** | 0.001112 | n/a | n/a |
| ***asy1/+*** | 0.000522 | 0.0007174 | n/a |
| ***asy1/+ cmt3*** | 0.001237 | 0.0123688 | 0.00123689 |

**Supplemental Table 16.** Genetic distances of *CTL3.2* in wild type, *cmt3*, *asy1/+* and *cmt3 asy1/+*. cM were calculated using the formula: cM = 100 × (1 – [1-2(*N_G_*+*N_R_*)/*N_T_*] ^½^), where *N_G_* is a number of green-alone fluorescent seeds, *N_R_* is a number of red-alone fluorescent seed and *N_T_* is the total number of seeds counted.

| Genotype | Green-alone | Red-alone | Both red and green | Neither red or green | Total | CTL3.2, cM | Average, CTL3.2, cM |
| --- | --- | --- | --- | --- | --- | --- | --- |
| Wild type | 165 | 181 | 1509 | 374 | 2229 | 16.96 |  |
| Wild type | 154 | 182 | 1478 | 364 | 2178 | 16.85 |  |
| Wild type | 188 | 166 | 1452 | 364 | 2170 | 17.92 |  |
| Wild type | 173 | 191 | 1487 | 403 | 2254 | 17.72 |  |
| Wild type | 182 | 145 | 1313 | 382 | 2022 | 17.75 |  |
| Wild type | 174 | 198 | 1527 | 370 | 2269 | 18.02 |  |
| Wild type | 175 | 186 | 1690 | 402 | 2453 | 16.00 |  |
| Wild type | 157 | 146 | 1416 | 356 | 2075 | 15.86 | 17.13 |
| *cmt3* | 153 | 165 | 1446 | 366 | 2130 | 16.25 |  |
| *cmt3* | 149 | 172 | 1370 | 378 | 2069 | 16.95 |  |
| *cmt3* | 130 | 185 | 1412 | 323 | 2050 | 16.77 |  |
| *cmt3* | 143 | 146 | 1370 | 357 | 2016 | 15.54 |  |
| *cmt3* | 162 | 159 | 1414 | 386 | 2121 | 16.49 |  |
| *cmt3* | 187 | 160 | 1604 | 384 | 2335 | 16.17 |  |
| *cmt3* | 207 | 174 | 1693 | 393 | 2467 | 16.87 |  |
| *cmt3* | 205 | 210 | 1703 | 469 | 2587 | 17.59 |  |
| *cmt3* | 181 | 180 | 1642 | 401 | 2404 | 16.35 | 16.55 |
| *asy1/+* | 188 | 239 | 1443 | 357 | 2227 | 21.48 |  |
| *asy1/+* | 210 | 196 | 1378 | 317 | 2101 | 21.67 |  |
| *asy1/+* | 197 | 186 | 1263 | 330 | 1976 | 21.75 |  |
| *asy1/+* | 229 | 214 | 1460 | 361 | 2264 | 21.98 |  |
| *asy1/+* | 228 | 209 | 1651 | 428 | 2516 | 19.21 |  |
| *asy1/+* | 206 | 193 | 1513 | 369 | 2281 | 19.37 |  |
| *asy1/+* | 222 | 175 | 1526 | 363 | 2286 | 19.21 |  |
| *asy1/+* | 251 | 169 | 1546 | 432 | 2398 | 19.40 |  |
| *asy1/+* | 165 | 147 | 1142 | 294 | 1748 | 19.81 |  |
| *asy1/+* | 168 | 179 | 1205 | 329 | 1881 | 20.56 |  |
| *asy1/+* | 157 | 150 | 1168 | 307 | 1782 | 19.04 | 20.32 |
| *cmt3 asy1/+* | 188 | 178 | 1435 | 382 | 2183 | 18.47 |  |
| *cmt3 asy1/+* | 205 | 184 | 1519 | 421 | 2329 | 18.39 |  |
| *cmt3 asy1/+* | 160 | 174 | 1167 | 284 | 1785 | 20.89 |  |
| *cmt3 asy1/+* | 171 | 180 | 1425 | 397 | 2173 | 17.72 |  |
| *cmt3 asy1/+* | 213 | 175 | 1499 | 402 | 2289 | 18.70 |  |
| *cmt3 asy1/+* | 169 | 172 | 1602 | 432 | 2375 | 15.57 |  |
| *cmt3 asy1/+* | 187 | 192 | 1693 | 449 | 2521 | 16.37 |  |
| *cmt3 asy1/+* | 219 | 193 | 1747 | 466 | 2625 | 17.17 |  |
| *cmt3 asy1/+* | 197 | 191 | 1663 | 459 | 2510 | 16.88 |  |
| *cmt3 asy1/+* | 191 | 173 | 1679 | 472 | 2515 | 15.71 | 17.59 |

**Supplemental Table 17.** Statistical significance (Mann-Whitney test) of genetic distances of *CTL3.2* in wild type, *cmt3*, *asy1/+* and *cmt3 asy1/+*.

|  | **Wild type** | ***cmt3*** | ***asy1/+*** |
| --- | --- | --- | --- |
| ***cmt3*** | 0.282499 | n/a | n/a |
| ***asy1/+*** | 0.001632 | 0.001176 | n/a |
| ***asy1/+ cmt3*** | 0.656688 | 0.282499 | 0.003280 |

**Supplemental Table 18.**
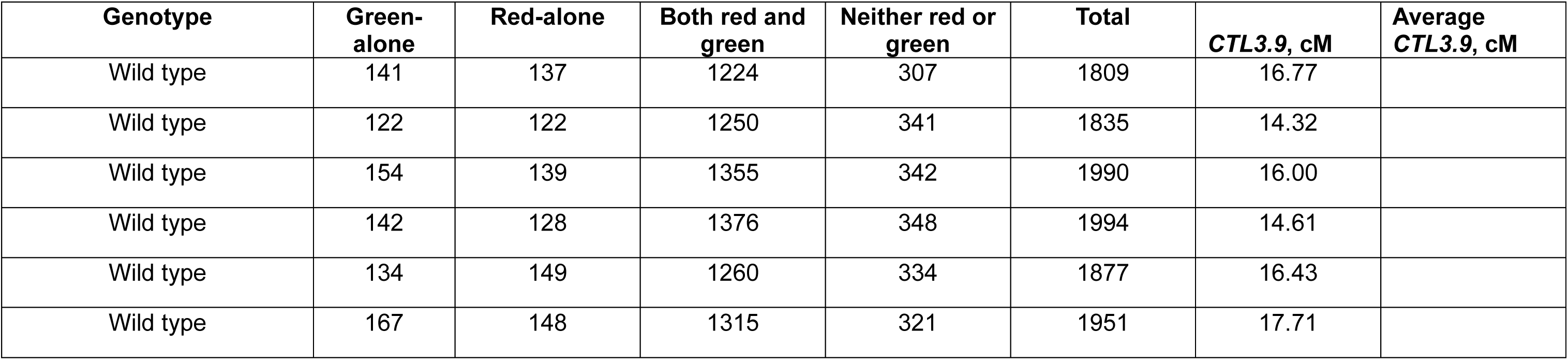
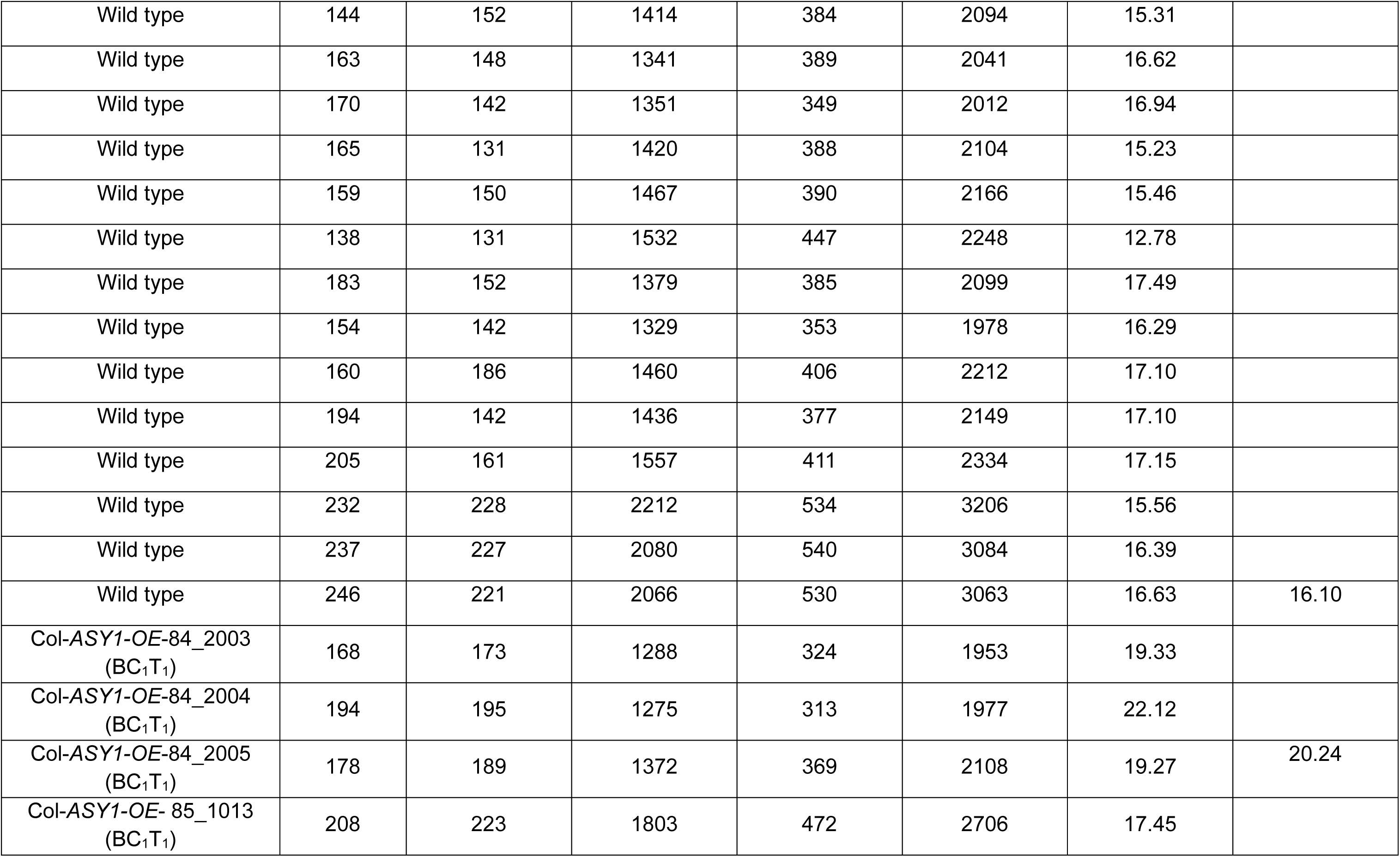
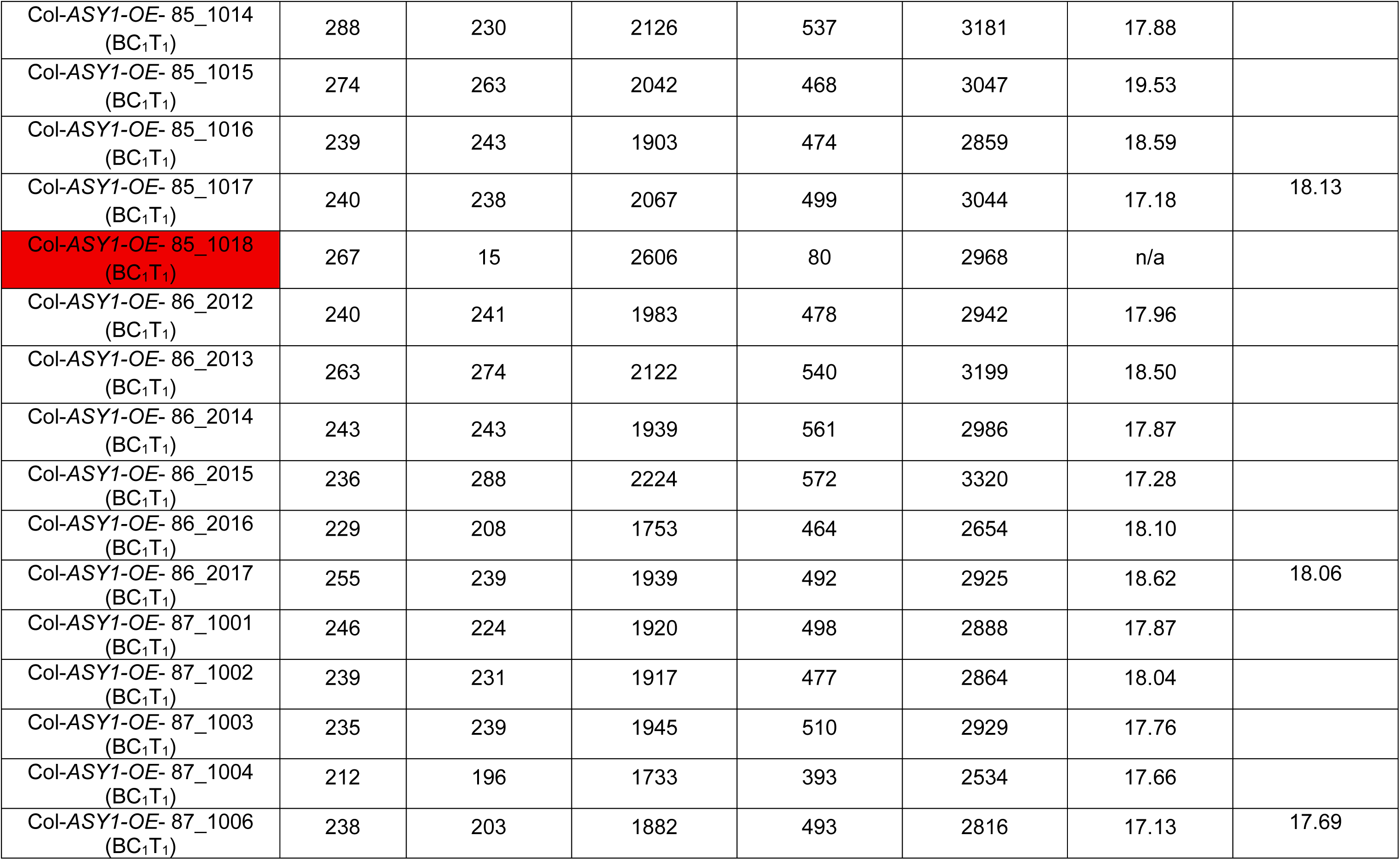
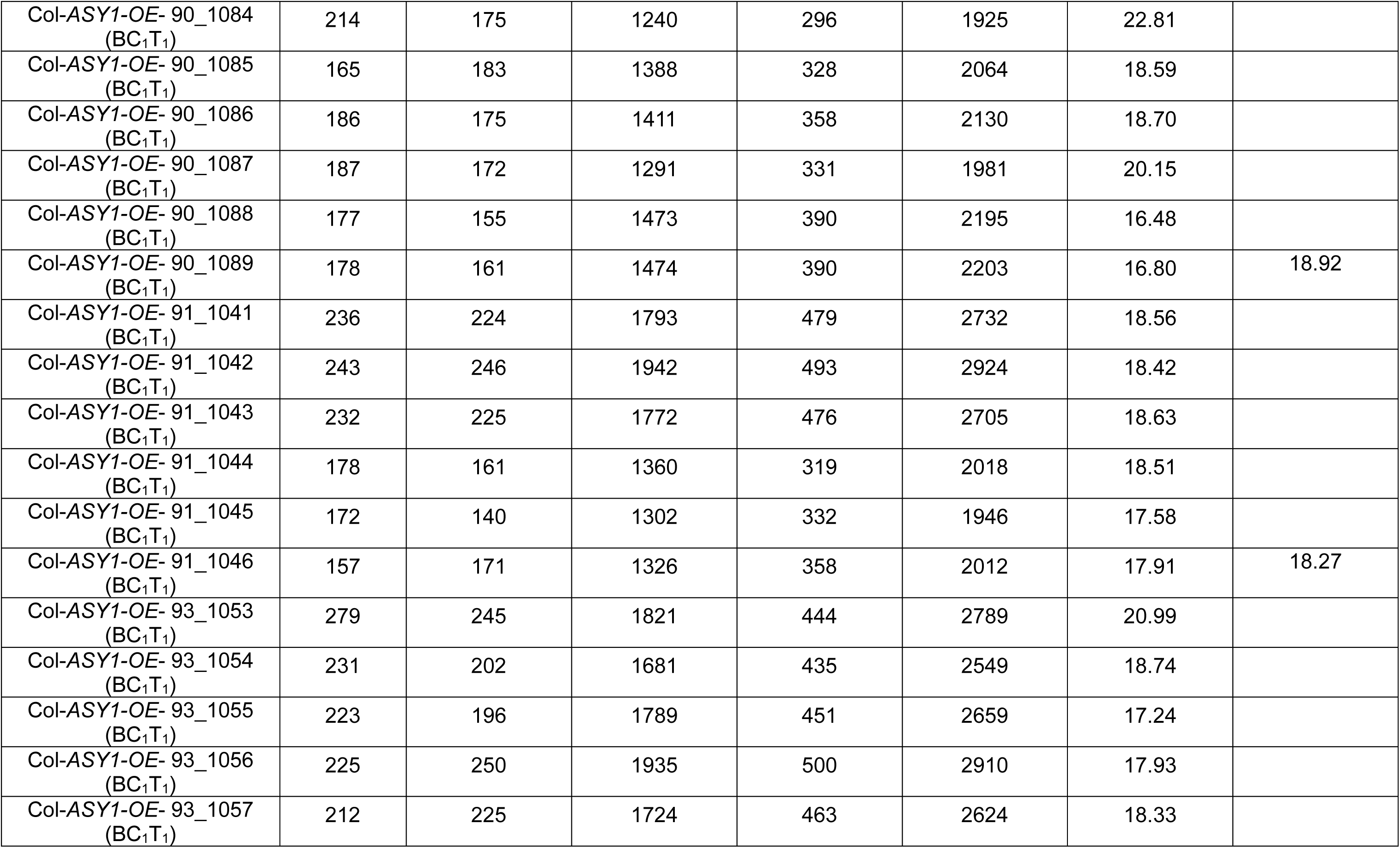
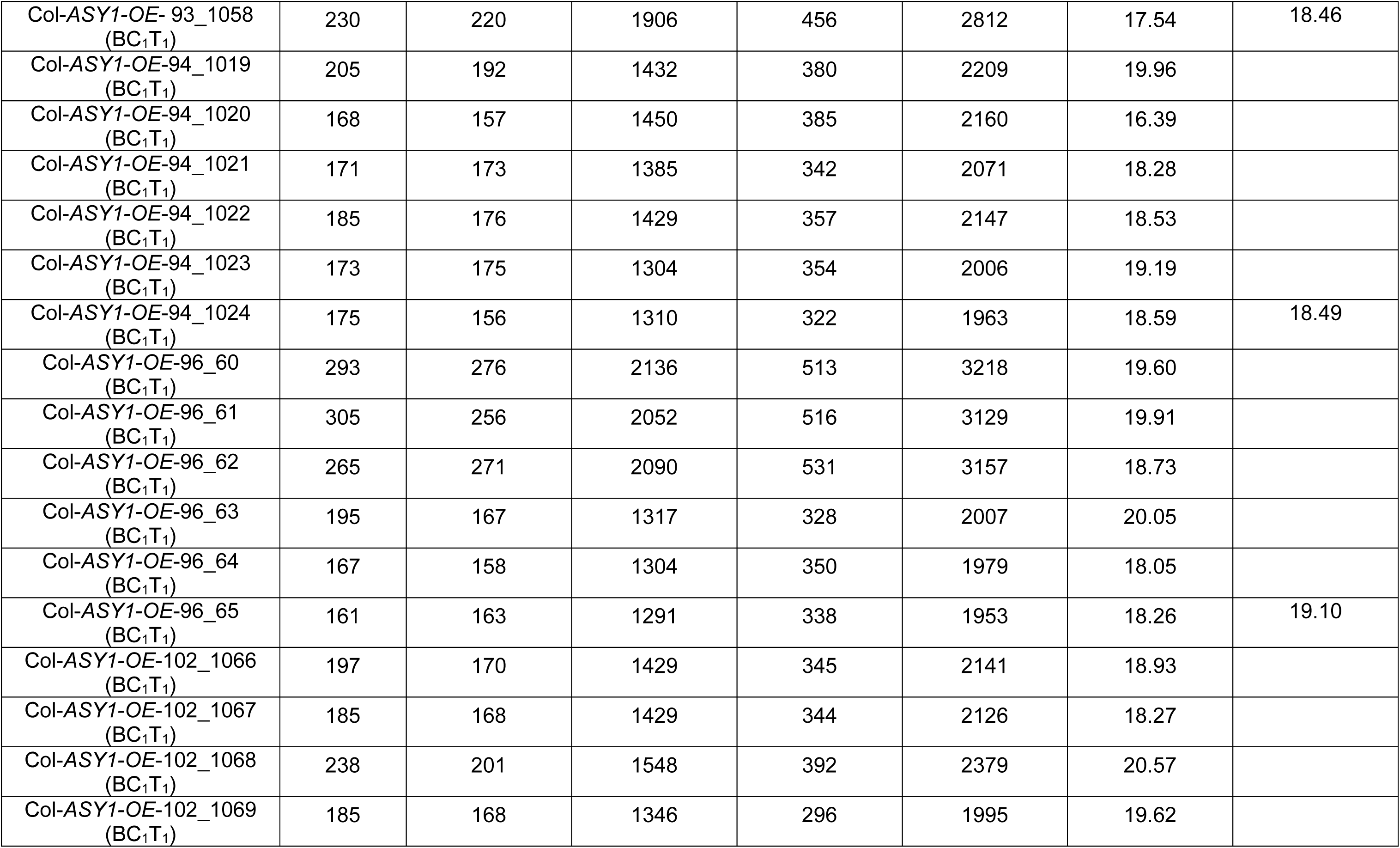
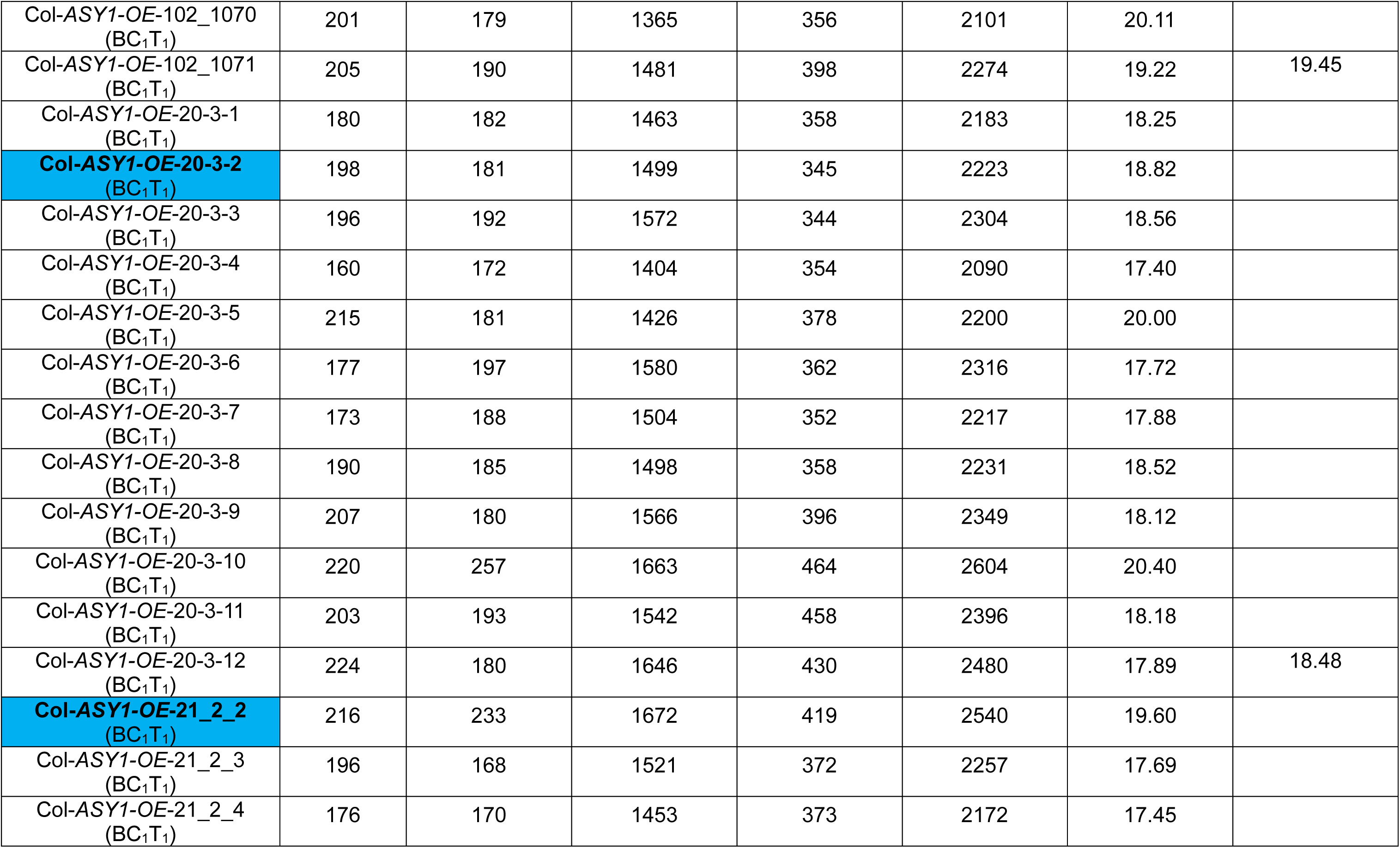
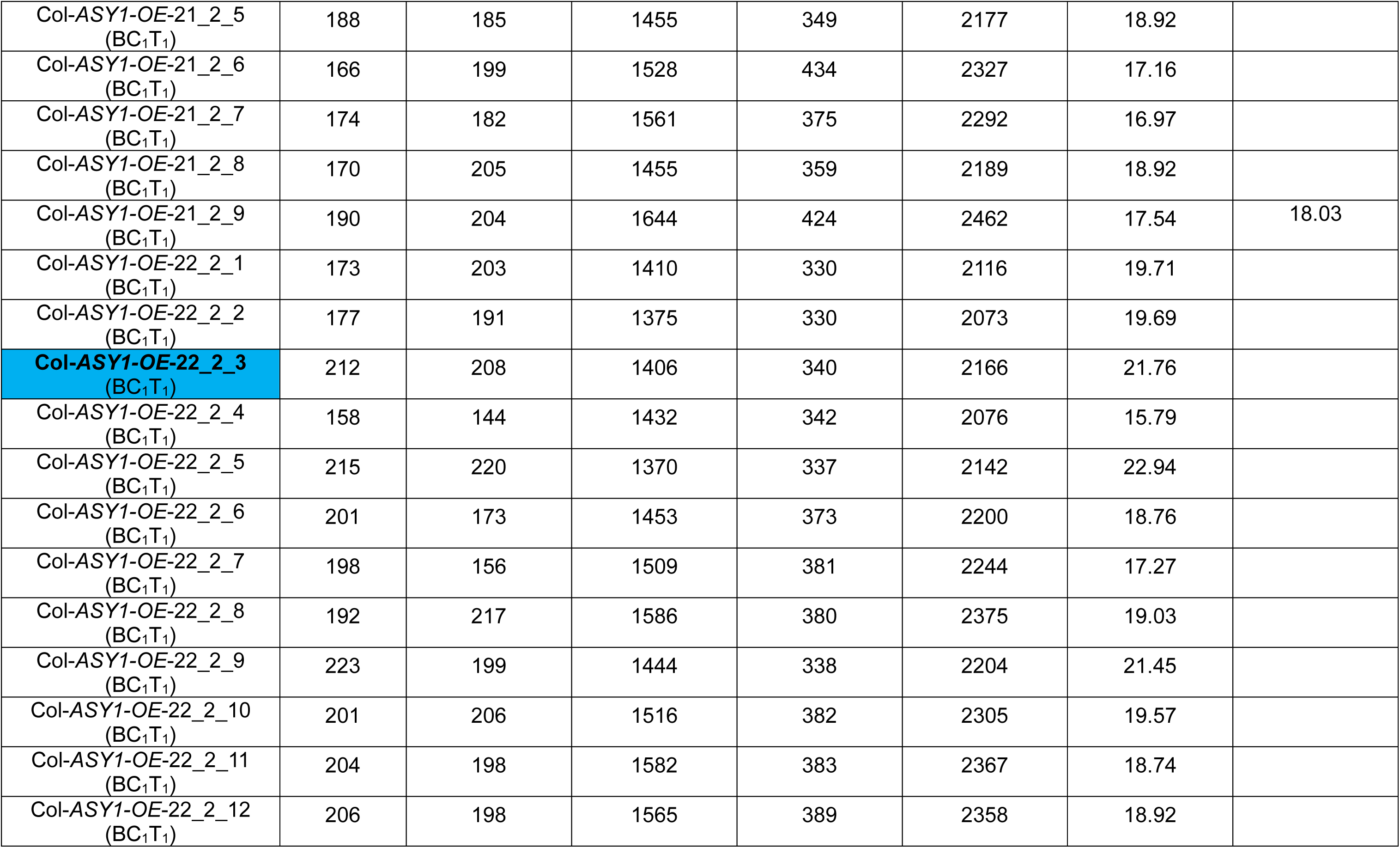
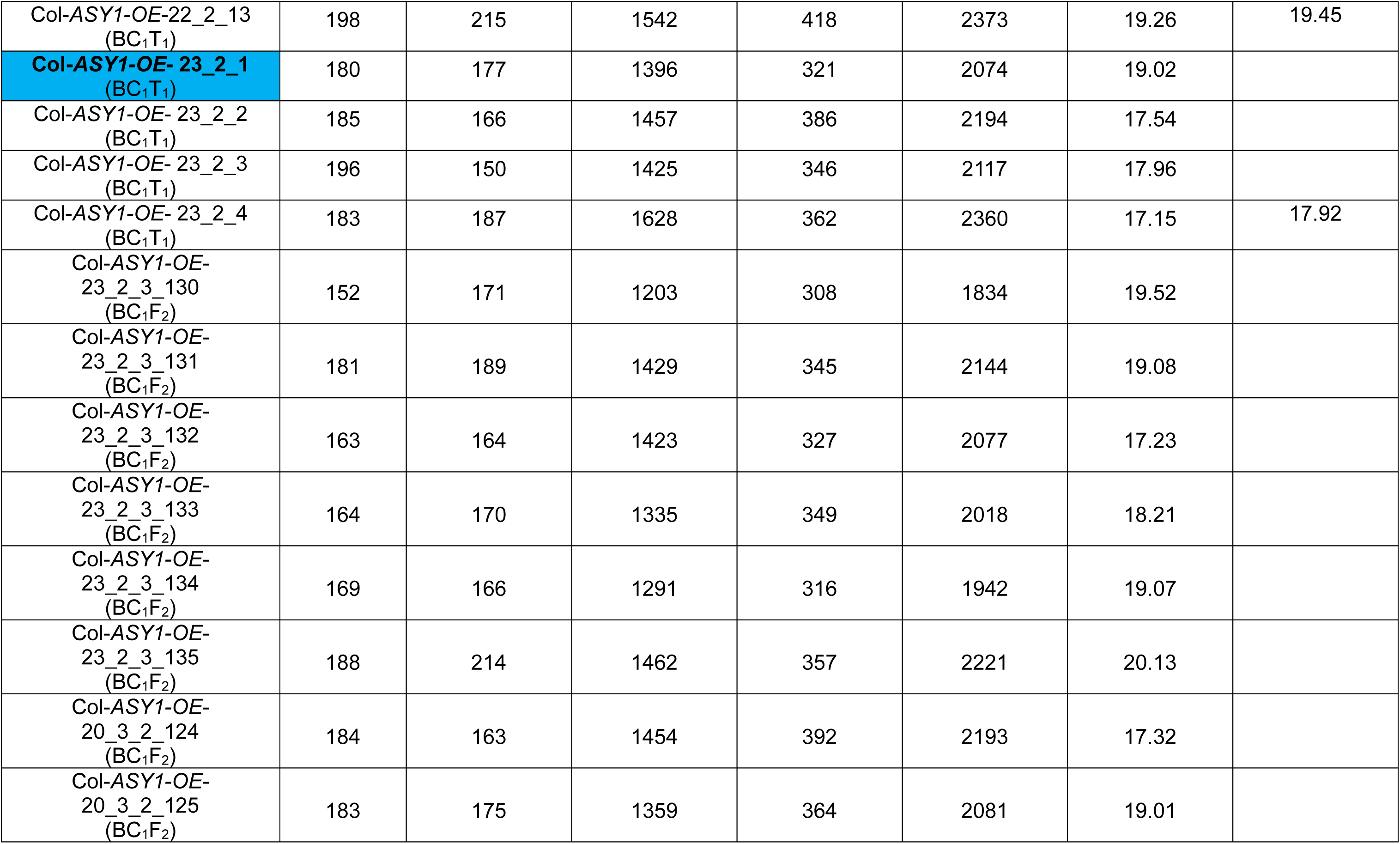
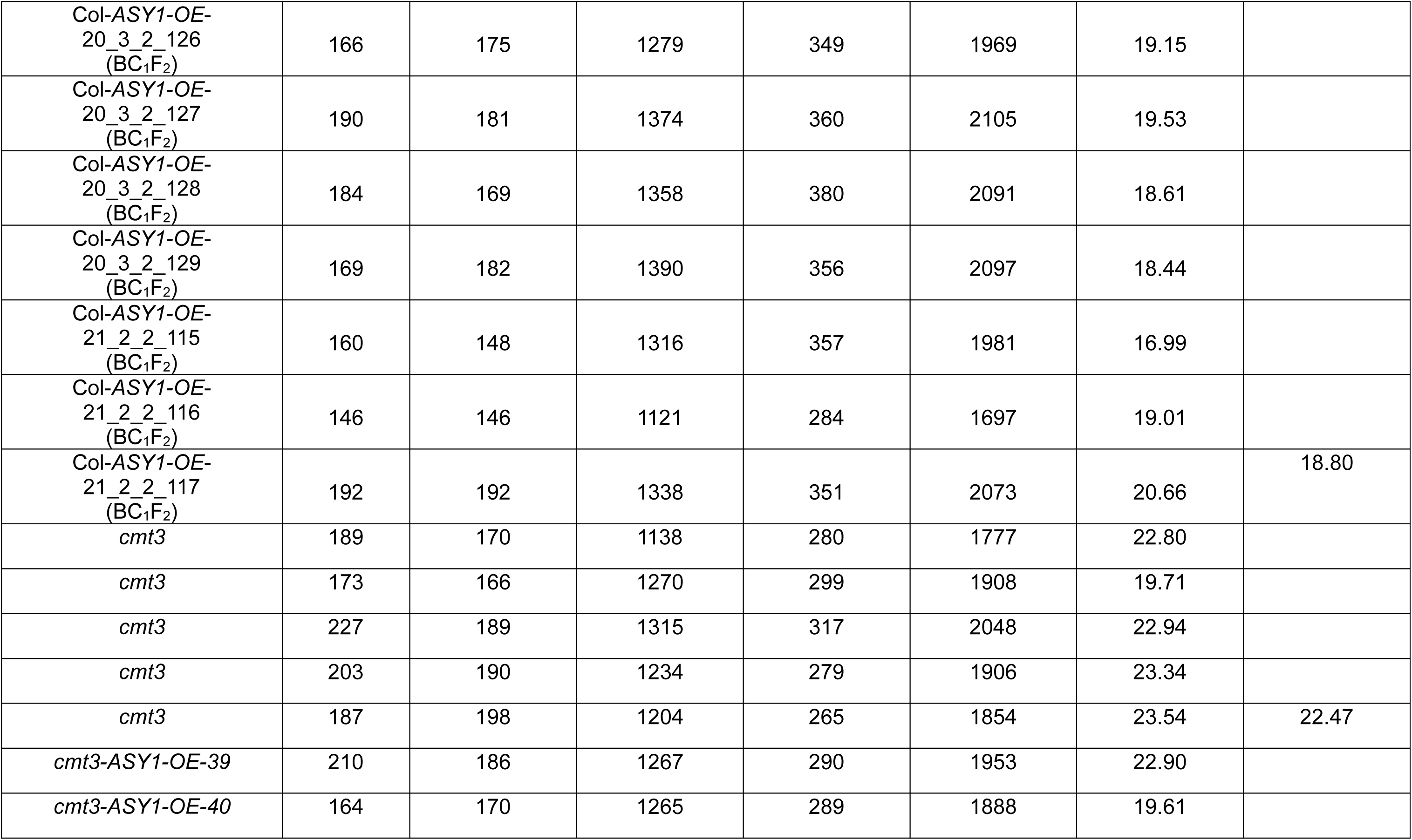
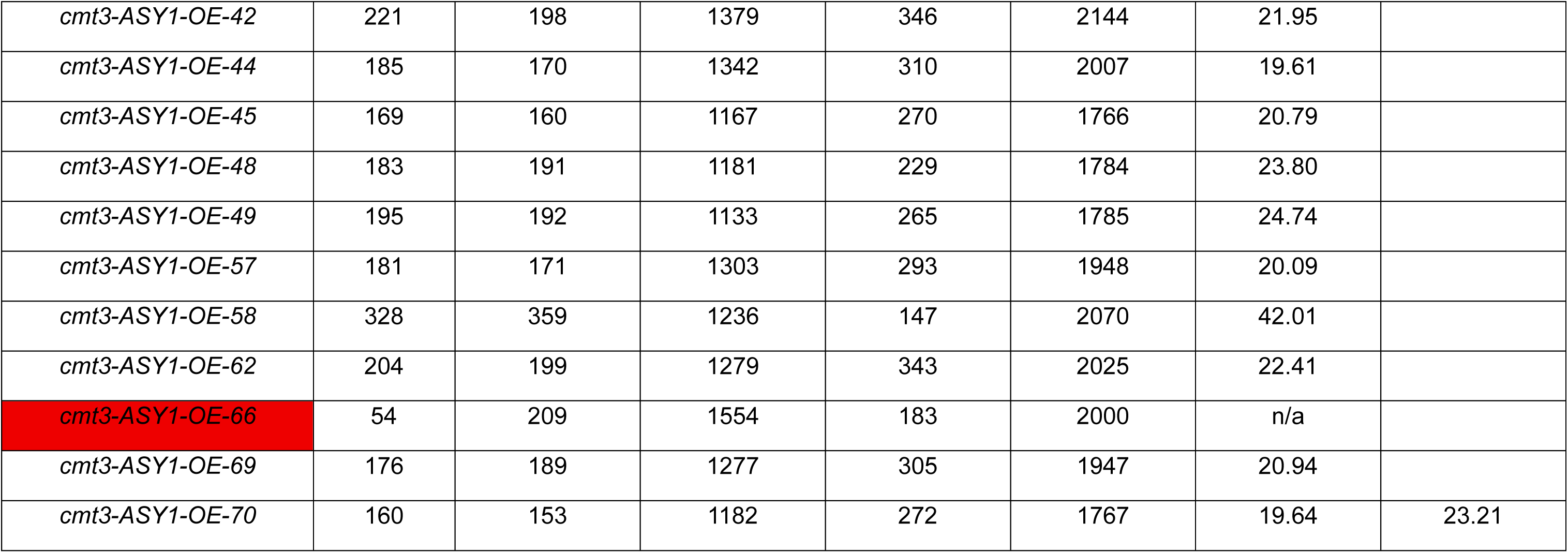
Genetic distances of *CTL3.9* in the wild type, *ASY1-OE* (BC_1_T_1_ and BC_1_F_2_), *cmt3*, and *cmt3 ASY1-OE* (T_1_). cM were calculated using the formula: cM = 100 × (1 – [1-2(*N_G_*+*N_R_*)/*N_T_*] ^½^), where *N_G_* is a number of green-alone fluorescent seeds, *N_R_* is a number of red-alone fluorescent seed and *N_T_* is the total number of seeds counted. Plants with segregation distortion of *eGFP* and *dsRed* are highlighted in red. BC_1_F_1_ individuals that were self-fertilised to generate BC_1_F_2_ are highlighted in blue.

**Supplemental Table 19.** Statistical significance (Mann-Whitney test) of genetic distances of *CTL3.9* in wild type, *ASY1-OE*, *cmt3*, and *cmt3 ASY1-OE*.

|  | **Wild type** | ***ASY1*-OE (BC<sub>1</sub>T<sub>1</sub>)** | ***ASY1*-OE (BC<sub>1</sub>F<sub>2</sub>)** | ***cmt3*** |
| --- | --- | --- | --- | --- |
| ***ASY1*-OE (BC<sub>1</sub>T<sub>1</sub>)** | 1.146*10<sup>-9</sup> | N/A | N/A | N/A |
| ***ASY1*-OE (BC<sub>1</sub>F<sub>2</sub>)** | 2.25*10<sup>-5</sup> | 0.692 | N/A | N/A |
| ***cmt3*** | 3.078*10<sup>-3</sup> | 2.629*10<sup>-3</sup> | 0.00672 | N/A |
| ***cmt3*-*ASY1*-OE (T<sub>1</sub>)** | 2.297*10<sup>-5</sup> | 1.306*10<sup>-5</sup> | 0.000417 | 0.692 |

**Supplemental table 20.** Distortion of segregation of *eGFP* and *dsRed* in the *CTL3.9* reporter line observed in two *ASY1-OE* plants (shaded in grey). Cells showing deviation from the expected ratios (Green/non Green and Red/non Red should equal 3, and Green/Red should equal 1) are shaded in red. A Col wild type plant with expected *eGFP* and *dsRed* segregation ratios is shown for comparison.

| Sample name | Green | Red | Both | Neither red nor green | Total | Green+Red/Total | Recombination frequency, cM | Green/non Green | Red/non Red | Green/Total | Red/Total | Green/Red |
| --- | --- | --- | --- | --- | --- | --- | --- | --- | --- | --- | --- | --- |
| Col-ASY1-OE_85_1018 | 267 | 15 | 2606 | 80 | 2968 | 0.10 | N/A | 30.24 | 7.55 | 0.09 | 0.01 | 17.80 |
| ASY1-OE-66, *cmt3* | 54 | 209 | 1554 | 183 | 2000 | 0.13 | N/A | 4.10 | 7.44 | 0.03 | 0.10 | 0.26 |
| Col wild type | 141 | 137 | 1224 | 307 | 1809 | 0.15 | 16.77 | 3.07 | 3.04 | 0.08 | 0.08 | 1.03 |

